# DOT1L limits regenerative activation of skeletal stromal progenitors

**DOI:** 10.64898/2026.04.06.716818

**Authors:** Marta Stetsiv, Drew Dauphinee, Sakinah Abdulsalam, Shagun Prabhu, Alexander Tress, Kerry Cobb, Archana Sanjay, Rosa M. Guzzo

**Affiliations:** Department of Neuroscience, School of Medicine, UConn Health; Department of Orthopaedic Surgery, School of Medicine, UConn Health; Computational Biology Core, UConn Health 263 Farmington Avenue, Farmington, CT, 06030 USA

**Keywords:** CXCL12-abundant reticular (CAR) cells, Dot1L histone methyltransferase, epigenetic regulation, intramembranous ossification, skeletal stem and progenitor cells

## Abstract

Adult bone marrow contains a heterogeneous network of skeletal stromal and progenitor cells (SSPC) that maintain bone homeostasis, support the hematopoietic niche, and drive the regenerative responses to injury. Despite their central role in bone regeneration, the mechanisms that regulate injury-induced SSPC activation and fate transitions remain incompletely understood. Here we identify Disruptor of telomeric silencing 1 like (DOT1L), the sole histone methyltransferase responsible for H3K79 methylation, as a critical regulator that restrains stromal activation. Using complementary genetic, pharmacologic, single-cell transcriptomic and injury models, we show that either Prrx1 lineage *Dot1L* haploinsufficiency or acute pharmacological Dot1L inhibition promote SSPC expansion, but through distinct programs: haploinsufficiency drives the expansion of Cxcl12+ CAR cells, whereas acute pharmacologic inhibition enriches fibroblastic-like stromal states. Single Cell Regulatory Network Inference and Clustering (SCENIC) analysis identifies DOT1L as a stabilizer of homeostatic marrow-supportive transcriptional programs, whose disruption facilitates transition to injury responsive stromal states. Partial loss of *Dot1L* in the Prrx1 lineage enhances injury-induced intramedullary mineralization *in vivo*. Collectively, these findings establish DOT1L as a gatekeeper of the stromal progenitor state that restrains lineage commitment and restrains the magnitude of the SSPC response following injury.

## INTRODUCTION

Significant efforts have been dedicated toward resolving the heterogeneity of the adult bone marrow stromal compartment, and its functional contributions to bone homeostasis and repair. Early foundational studies established that perisinusoidal stromal cells expressing the leptin receptor (LEPR) represent the principal source of osteoblasts and adipocytes in adult bone marrow. These cells remain largely quiescent under homeostatic conditions but become activated following skeletal injury.^1, 2^ Closely related CXCL12-abundant reticular (CAR) cells, located in perisinusoidal niches, are best known for producing critical niche-supportive factors such as CXCL12 and Stem Cell Factor (SCF) that sustain hematopoietic stem and progenitor cells.^3, 4, 5^ Genetic cell-fate tracking and single cell-RNA sequencing (scRNA-seq) studies have further revealed that osteogenic support within the marrow arises from multiple specialized CAR subsets with distinct activation states and lineage biases. ^6, 7, 8, 9, 10^ Adipo-CAR cells (*Cxcl12+, LepR+, Alpl-*) localized to areas around sinusoidal vessels in the central marrow, and osteo-CAR cells (*Cxcl12+*, *LepR-*, *Alpl+*) localized to arterioles in close proximity to bone.^6, 11^ Together, these studies indicate that CAR heterogeneity reflects a spectrum of lineage priming rather than a single uniform progenitor state. Consistent with this framework, Liao and colleagues identified a distinct Adipoq⁺ Osx⁺ endosteal progenitor population in adult bone marrow that contributes to both osteogenic and adipogenic lineages.^12^

Bone injury models have demonstrated that marrow CAR populations undergo rapid activation, proliferative expansion, and transcriptional reprogramming following bone damage, enabling their migration toward injury sites and differentiation into osteolineage cells.^13^ Together, these findings establish perisinusoidal CXCL12⁺ stromal cells as a major source of injury-responsive skeletal stromal and progenitor cells (SSPCs) in adult bone marrow. Significantly, CXCL12+ stromal populations retain multilineage potential. Effective skeletal repair depends not only on their activation, but also on appropriate regulation of lineage engagement during regeneration.^14^ Important questions remain regarding the mechanisms which coordinate the transition of these diverse CAR populations from quiescent niche supporting to lineage-committed states during injury.

Although classical signaling pathways including WNT, BMP and NOTCH have established roles in regulating SSPC activation and lineage commitment,^13, 15, 16, 17, 18, 19^ little is known about the epigenetic mechanisms that establish or restrict progenitor competence during adult skeletal regeneration. Increasing evidence suggests that chromatin modifiers function in a context-dependent manner to modulate progenitor quiescence, activation, lineage permissiveness, and cellular plasticity in response to developmental and environmental cues, rather than functioning as universal regulators of differentiation. ^20, 21, 22^ Whether specific chromatin modifiers actively gate the transition of CAR cells from a quiescent, niche-supportive state to an activated, lineage-committed state remains unknown. Defining how these epigenetic mechanisms regulate CAR cell transitions is essential to improve the fundamental understanding of native skeletal maintenance and repair processes.

Disruptor of Telomeric Silencing 1 Like (DOT1L) is the sole histone methyltransferase responsible for methylating lysine 79 in histone 3 (H3K79me1/2/3).^23^ Accumulating evidence suggests that DOT1L-dependent H3K79 methylation functions as a context-dependent modulator of lineage specific transcriptional networks, promoting or constraining gene expression depending on the cellular states.^24, 25, 26, 27, 28^ The loss of DOT1L function accelerates differentiation trajectories and disrupts the balance between proliferation and maturation.^25^ Studies across neural, hematopoietic and immune tissues revealed that the loss of DOT1L expression or inhibition of its enzymatic activity biases progenitors towards more mature, differentiated cell states, indicating that DOT1L functions in gating progenitor cell differentiation^29, 30, 31, 32, 33 28, 34^ Within the skeletal system, DOT1L has been established as a critical regulator of skeletal development, endochondral ossification,^35^ and articular cartilage homeostasis.^36, 37, 38^ However, regeneration requires the activation and reprogramming of quiescent adult skeletal progenitors in response to injury, raising the possibility that DOT1L serves distinct functions in developmental and regenerative contexts.

Here, we combined *Dot1L* genetic perturbation and pharmacologic inhibition of DOT1L methyltransferase activity with single-cell transcriptomics, trajectory inference, gene regulatory network reconstruction and functional analyses to define the role of DOT1L in bone marrow SSPCs during adult skeletal maintenance and induction of injury response programs. We demonstrate that DOT1L acts as an epigenetic regulator of bone marrow stromal cell plasticity by maintaining CAR cell identity and restraining transitions toward activated lineage-committed states. Reduced DOT1L activity remodels transcriptional regulatory networks, expands injury-responsive stromal progenitor populations, and enhances intramedullary bone formation following mechanical marrow injury, identifying DOT1L as a critical regulator of marrow SSPCs.

## RESULTS

### Dot1L exhibits widespread expression throughout the bone marrow stromal compartment

To determine whether DOT1L is positioned to regulate bone marrow stromal progenitor populations, we examined its expression across an integrated scRNA-seq dataset comprising murine bone marrow cells from multiple experimental and physiological conditions.^39^ This harmonized dataset encompasses diverse marrow cell types, including stromal, hematopoietic, endothelial, and other niche-associated populations, and is organized into transcriptionally distinct clusters annotated based on established lineage and niche marker expression (**Supplemental Figure 1a**). Projection of *Dot1L* gene expression onto this clustered dataset showed widespread transcript detection across bone marrow clusters expressing canonical markers of stromal progenitor states (**Supplemental Figure 1b**).

To more precisely define *Dot1L* expression within stromal subsets relevant to bone regeneration, we analyzed a published scRNA-seq dataset generated from CXCL12-enriched bone marrow stromal cells isolated from mouse femurs (GSE136979).^13^ Unsupervised clustering revealed transcriptionally heterogeneous populations within the CXCL12*+* marrow niche, including reticular and stromal populations marked by canonical niche-associated genes such as *Cxcl12*, *Adipoq, Lepr,* and *Pdgfra* (**Figure 1a,b,d**). A smaller subset of this population was identified as pre-osteoblasts, marked by expression of *Alpl, Col1a1, Wif1, Tnc, Ly6a,* and *Thy1* (**Figure 1a**). Additional non-stromal clusters corresponding to endothelial, immune, and erythroid populations were also identified. *Dot1L* transcripts were broadly detected throughout the reticular and stromal populations within this marrow microenvironment (**Figure 1c**). Overlay of stromal lineage marker expression further confirmed expression of *Dot1L* within a CXCL12+ subset expressing *Prrx1, Pdgfra, Lepr, Adipoq,* and *Col1a1* (**Figure 1d**), indicating that *Dot1L* is expressed across marrow-resident mesenchymal and stromal progenitor compartments relevant to skeletal repair. Consistent with emerging models emphasizing epigenetic regulators as integrators of progenitor quiescence, activation, and injury-induced fate transitions in bone,^20, 22^ gene expression analyses of chromatin modifying enzymes implicated in osteogenic regulation showed that DOT1L is part of a wider chromatin-regulatory landscape within the broad stomal populations (**Supplemental Figure 1c**) and more specifically in the CXCL12^+^ marrow niche (**Supplemental Figure 1d**). Given its unique role as the sole H3K79 methyltransferase, these findings formed the rationale for the present study, which tested whether DOT1L functions as a non-redundant regulator of adult bone marrow SSPC identity, activation and state transitions, and injury-induced osteogenesis.

**Figure 1.**
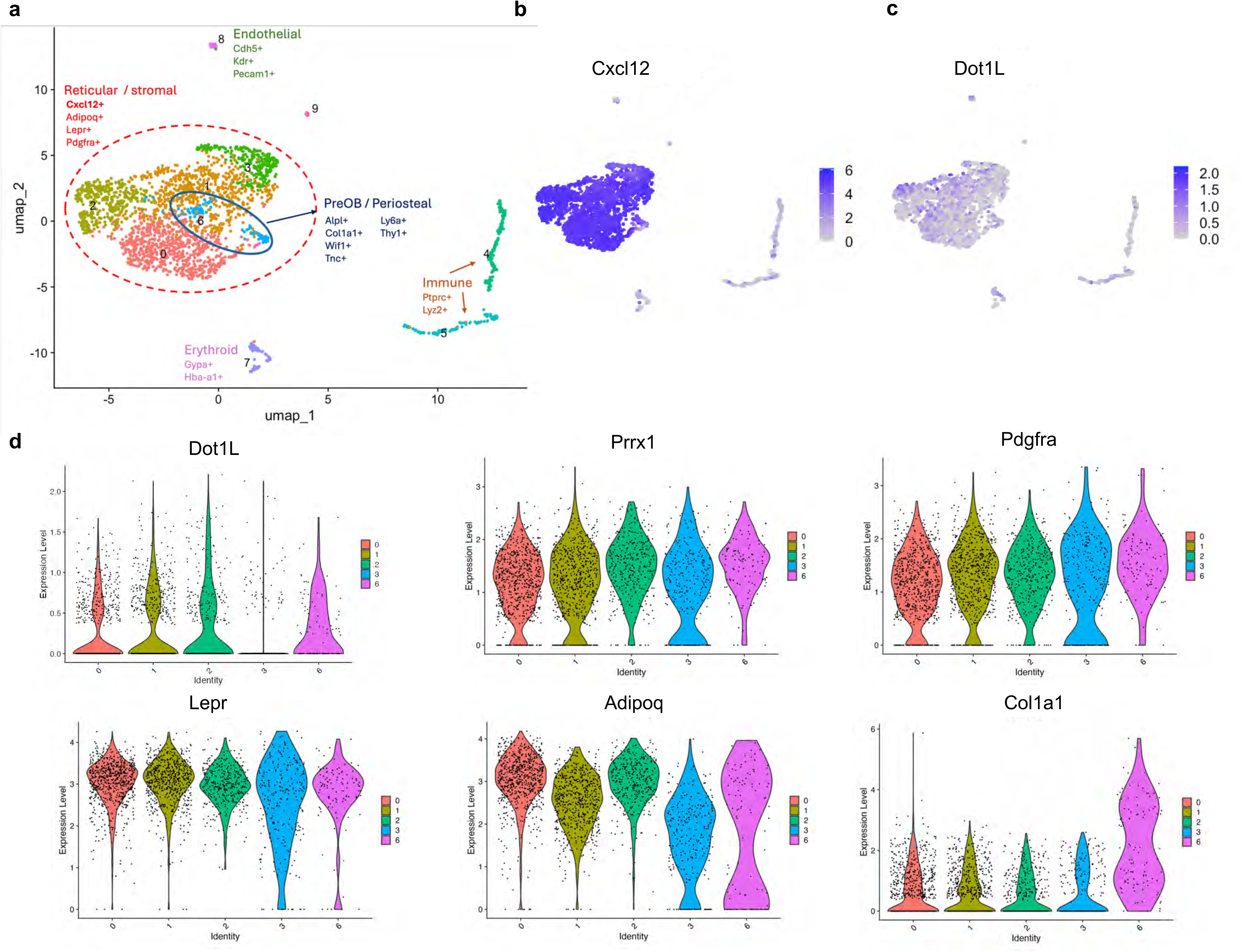
*Dot1L* expression within subsets of mouse CXCL12^+^ bone marrow cells. (**a**) Uniform manifold approximation (UMAP) plot of unsupervised clustering revealed transcriptionally distinct sub-populations of CXCL12^+^ Lineage^−^ mouse bone marrow cells (GSE136979)^13^. The population was largely defined by reticular stromal properties including the expressions of *Cxcl12*, *Adipoq*, *Lepr*, and *Pdgfra* (dashed red line outlining clusters 0, 1, 2, 3, and 6). A subset of stromal cells displayed a transcriptional profile consistent with pre-osteoblast/periosteal lineage identity (blue outline, cluster 6). Additional clusters correspond to endothelial, immune, and erythroid populations (clusters 4, 5, 7, 8 and 9). (**b, c**) UMAP feature plots using color to indicate gene expression (log CPM) of *Cxcl12 and Dot1L.* (**d**) Violin plots demonstrate the enrichment of *Dot1L* within stromal and osteoprogenitor subsets expressing *Prrx1, Pdgfra, Lepr, AdipoQ* and *Col1a1*.

### Genetic deletion of Dot1L by Prrx1Cre expanded the bone marrow stromal compartment and remodeled transcriptional regulatory programs under homeostatic conditions

We employed single cell RNA sequencing (scRNA-seq) to comprehensively investigate the role of DOT1L in the steady-state regulation of the bone marrow stromal compartment. For these studies, we used the Prrx1-Cre driver to delete *Dot1L* within the limb mesenchymal lineage, resulting in loss of DOT1L expression within skeletal tissues and bone marrow stromal cells.^35^ The CD45^−^CD31^−^Ter119^−^ (Lineage^−^) bone marrow cells harvested from the femurs and tibias of control Dot1L^fl/fl^ and Dot1L^fl/fl^:Prrx1Cre adult mice were enriched by FACS (**Supplemental Figures 2a-c**), and analyzed in downstream scRNA sequencing. After quality filtering (**Supplemental Figures 2d-f**), and removal of residual *Ptprc^+^*(CD45^+^) expressing cells, 3,945 Dot1L^fl/fl^ and 3,192 Dot1L^fl/fl^:Prrx1Cre high quality cells were recovered for analysis (**Supplemental Figure 3a,c**). Unsupervised clustering revealed 12 molecularly distinct cell populations within each sample (**Figure 2a** and **Supplemental Figure 3c**). Cell type identities were assigned to each cluster based on expression of established marker genes identified within the top differentially expressed, together with their overall abundance in each cluster (**Supplemental Figure 3b**). The clusters represented B cells (clusters 0, 2, 5, 6), granulocytes/neutrophils (clusters 1, 8, 10), erythroid cells (clusters 3, 7), and macrophages (clusters 4, 11). The frequency of hematopoietic cells associated with immature states (i.e. early erythroid cells, immature neutrophils, pre-B cells) was elevated in the Dot1L^fl/fl^:Prrx sample, along with concomitant reduction in terminally differentiated immune populations (i.e. mature neutrophils, mature B cells) relative to the control sample (**Figure 2a,b** and **Supplemental Figure 3c**), indicative of altered regulation within the hematopoietic niche. Given that Prrx1Cre is not active in these populations,^3, 5^ these observations likely reflect dysregulation through non-cell autonomous mechanisms.

**Figure 2.**
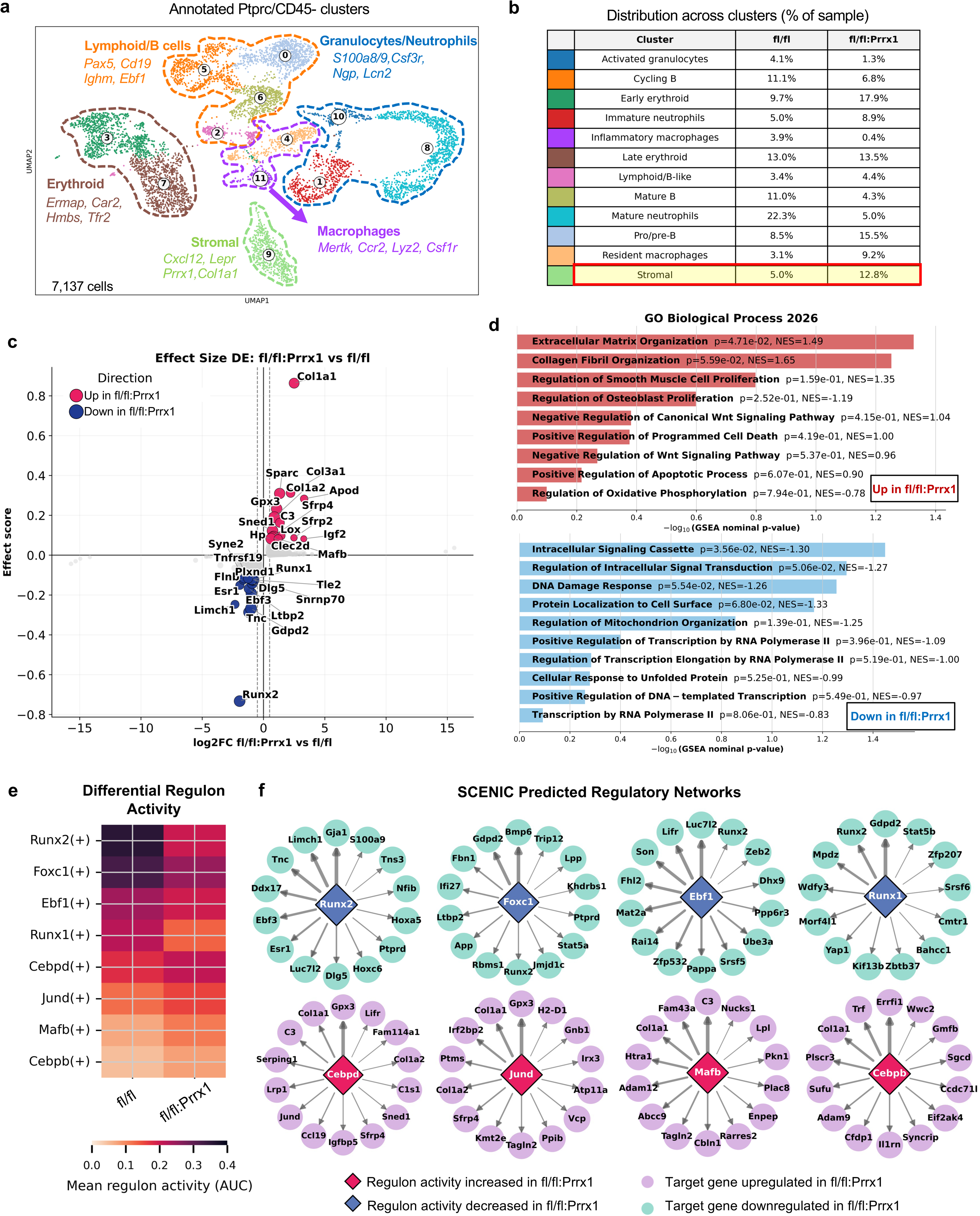
Single cell analysis of Lineage^−^ cells from bone marrow of Dot1L^fl/fl^ control and Dot1L^fl/fl^:Prrx1Cre mice. (**a)** UMAP plot showing annotated cell clusters from the analyses of 3,945 Dot1L^fl/fl^ cells and 3,192 Dot1L^fl/fl^:Prrx1Cre cells. (**b**) The percentage of cells within each cell cluster is shown for each sample. The frequency of cells in the stromal cluster are highlighted for each condition. (**c**) Differential gene expression within the stromal cluster ranked by effect score; top 15 up-regulated genes (magenta) and top 15 down-regulated genes (navy blue) in Dot1L^fl/fl^:Prrx1Cre cells are annotated. (**d**) Gene ontology (Biological Process 2026) analysis of stromal cluster pre-ranked genes showing the top enriched and attenuated processes in Dot1L^fl/fl^:Prrx1Cre. (**e**) Heatmap displays mean AUCell (AUC) regulon activity of select key transcription factor regulons with differential activity between Dot1L^fl/fl^:Prrx1Cre and Dot1L^fl/fl^ control cells. (**f**) Gene regulatory networks for individual regulons inferred by SCENIC show the top 14 targets ranked by regulon-target weight, with strength proportional to edge thickness. Diamond nodes represent regulons and circular nodes represent target genes.

Next, we assessed the impact of Prrx1Cre-mediated *Dot1L* deletion on the abundance and transcriptional profiles of cells within the stromal compartment (cluster 9) defined by expression of *Lepr, Cxcl12, Col1a1* and *Prrx1* itself (**Figure 2a** and **Supplemental Figure 3b**). This cluster exhibited a ∼2.5 fold expansion within the Dot1L^fl/fl^:Prrx1 sample as compared to the Dot1L^fl/fl^ control (**Figure 2b**). Effect size differential expression analysis revealed transcriptional changes indicative of stromal cell activation, loss of niche supportive stromal cell state, and extracellular matrix (ECM) remodeling in the Dot1L^fl/fl^:Prrx1Cre mice (**Figure 2c**). The most upregulated genes included ECM components and remodeling genes, such as *Col1a1, Col1a2, Col3a1, Sparc, Lox*, and *Sned1*. Additional increases in *Igf2, Sfrp2, Sfrp4, Apod, C3, Hp, Gpx3*, and *Clec2d* suggested activation of anabolic growth factor signaling, modulation of WNT signaling, oxidative stress responses, and broader stromal activation programs. In contrast, genes reduced in Dot1L^fl/fl^:Prrx1Cre were enriched for regulators of marrow stromal cell identity, skeletal homeostasis, and progenitor maintenance. These included the CAR-associated transcription factor *Ebf3*, the cytoskeletal regulator *Limch1*, the estrogen receptor *Esr1*, and additional genes involved in stromal organization and cell-cell interactions including *Runx1, Tek (Tie2), Plxnd1, Ltbp2, Dlg5, Flnb, Syne2,* and *Tnfrsf19*. Interestingly, expression of the osteogenic master regulator *Runx2* was also reduced despite the increased expression of ECM genes, suggesting that stromal cells were not undergoing overt osteogenic differentiation at baseline. Simultaneous loss of CAR-associated identity together with increased expression of ECM and activation-associated genes supports a transition toward an activated, matrix remodeling stromal state.

Gene ontology enrichment analysis (GSEA) was performed using pre-ranked genes upregulated in bone marrow stromal cells from Dot1L^fl/fl^:Prrx1Cre mice relative to Dot1L^fl/fl^ controls (**Figure 2d**). ECM organization (NES = 1.49) and collagen fibril organization (NES = 1.65) emerged as the dominant biological processes enriched in Dot1L^fl/fl^:Prrx1Cre mice, indicating broad activation of ECM production and remodeling programs. Consistent with the differential expression analysis, these pathways were driven by increased expression of multiple collagen genes (*Col1a1, Col1a2, Col3a1, Col6a3, Col6a5, Col18a1*, *Col26a1*), ECM structural and maturation genes (*Sparc*, *Lox*), and ECM remodeling enzymes (*Mmp2, Mmp28, Adamts1, Adamts5, Adamts15, Timp3*, *Timp4)*. Additional positively enriched pathways included regulation of osteoblast proliferation, negative regulation of canonical WNT signaling, supported by the differential expression of WNT pathway components and modulators including *Wnt5a, Wnt11, Fzd1, Sfrp2*, and *Sfrp4*. Collectively, these findings indicate that loss of *Dot1L* activates ECM remodeling and developmental signaling programs in the steady-state marrow, independent of injury. Conversely, pathways negatively enriched in Dot1L^fl/fl^:Prrx1Cre were associated with intracellular signaling and transcriptional regulation, including DNA damage response, and multiple RNA polymerase II-dependent transcriptional processes (**Figure 2d**). These pathway-level changes are consistent with the reduced expression of genes involved in marrow stromal cell identity and homeostasis, including *Ebf3, Limch1, Esr1, Runx1, Tek, Plxnd1*, and *Ltbp2*.

To identify the gene regulatory networks altered by loss of *Dot1L* we performed SCENIC regulon analysis (**Figures 2e,f**). In agreement with the differential expression and pathway enrichment analyses, transcription factor regulons exhibiting increased activity in Dot1L^fl/fl^:Prrx1Cre stromal cells, including *Cebpb*, *Cebpd*, *Jund, and Mafb*, were linked to transcriptional program governing stress adaptation, stromal cell activation, and tissue remodeling. Interestingly, these regulons converged on targets associated with ECM remodeling, stromal activation, and stress adaptation, including *Col1a1, Col1a2, Gpx3, Sfrp4, Lifr,* and *Tagln2*. In contrast, transcription factors associated with marrow stromal identity, niche-supportive functions, skeletal lineage specification, and osteogenic competence (*Foxc1, Ebf1, Runx2,* and *Runx1*) showed reduced regulon activity in Dot1L^fl/fl^:Prrx1Cre cells. Specifically, regulatory network inference analysis showed that TFs with lower activity in *Dot1L* deficient cells target stromal identity maintenance factors, including *Ebf3* and *Limch1*, and osteogenic factors *Bmp6*, *Tnc, Runx2*, and *Yap1*, consistent with DE analysis showing downregulation of these genes in Dot1L^fl/fl^:Prrx1Cre cells. Together, these findings indicate that *Dot1L* deficiency reprograms the stromal transcriptional network by suppressing stromal identity-maintaining programs while activating matrix remodeling pathways.

### Dot1L loss in the Prrx1 lineage promotes bone marrow stromal progenitor expansion and cell cycle entry in vivo

Transcriptomic analyses indicated the transition of DOT1L-deficient cells from homeostatic marrow stromal maintenance programs toward activated remodeling associated regulatory networks. Thus, we assessed whether Prrx1 driven loss of *Dot1L* modulates the proportion of cycling cells within the broader Lineage^−^ population. Cycling cells were detected primarily in myeloid (clusters 1, 8), erythroid (clusters 3, 7) and lymphoid lineages (cluster 5), with little to no detection of cycling stromal cells (cluster 9), indicative of their quiescence under steady-state conditions. Across the entire Lineage^−^ population, 25.6% of Dot1L^fl/fl^:Prrx1Cre cells were within S phase as compared to only 18.3% of Dot1L^fl/fl^ control cells (**Figure 3b**). Increased cycling of Lineage-cells from Dot1L^fl/fl^:Prrx1Cre mice was further validated by pulsing a separate cohort of adult Dot1L^fl/fl^:Prrx1Cre and Prrx1Cre control mice with EdU. Bone marrow cells were harvested directly and analyzed by flow cytometry (gating strategy in **Supplemental Figure 4**) without intervening culture, allowing assessment of proliferation and progenitor abundance under physiological conditions. Quantitative flow cytometric profiling confirmed a 1.5-fold increase in Lineage^−^ cells in the bone marrow of Dot1L^fl/fl^:Prrx1 mice compared with wild-type Prrx1-Cre (*p* = 0.002) (**Figures 3c,d**). Cell-cycle analysis through combined DNA content by Hoechst and EdU incorporation measurements showed a significantly increased proportion of cells entering S-phase in Dot1L^fl/fl^:Prrx1Cre compared to Prrx1-Cre controls (*p = 0.0002*), along with corresponding decrease in G0/G1 cells in Dot1L^fl/fl^:Prrx1Cre mice (*p = 0.0003*) (**Figures 3e,f**). Additionally, colony forming unit (CFU) assays showed increased clonogenic capacity in bone marrow mesenchymal cells harvested from Dot1L^fl/fl^:Prrx1 female and male mice relative to controls (**Figures 3g,h**). In parallel, quantitative assays using colorimetric cell counting kit-8 (CCK-8) confirmed that BMSCs from Dot1L^fl/fl^:Prrx1Cre mice of either sex retained a marked and highly significant increase in proliferative activity as compared to control Dot1L^fl/fl^ samples (**Figures 3h,j**). Together, these findings demonstrate that loss of *Dot1L* enhances the intrinsic clonal output and proliferative capacity of adult marrow stromal progenitors, consistent with an activated stromal state. Although cell cycle enrichment within the steady-state stromal compartment was limited by scRNAseq, increased proliferation across the broader Lineage-population suggests that reduced DOT1L activity may also influence neighboring populations through remodeling of the niche microenvironment.

**Figure 3.**
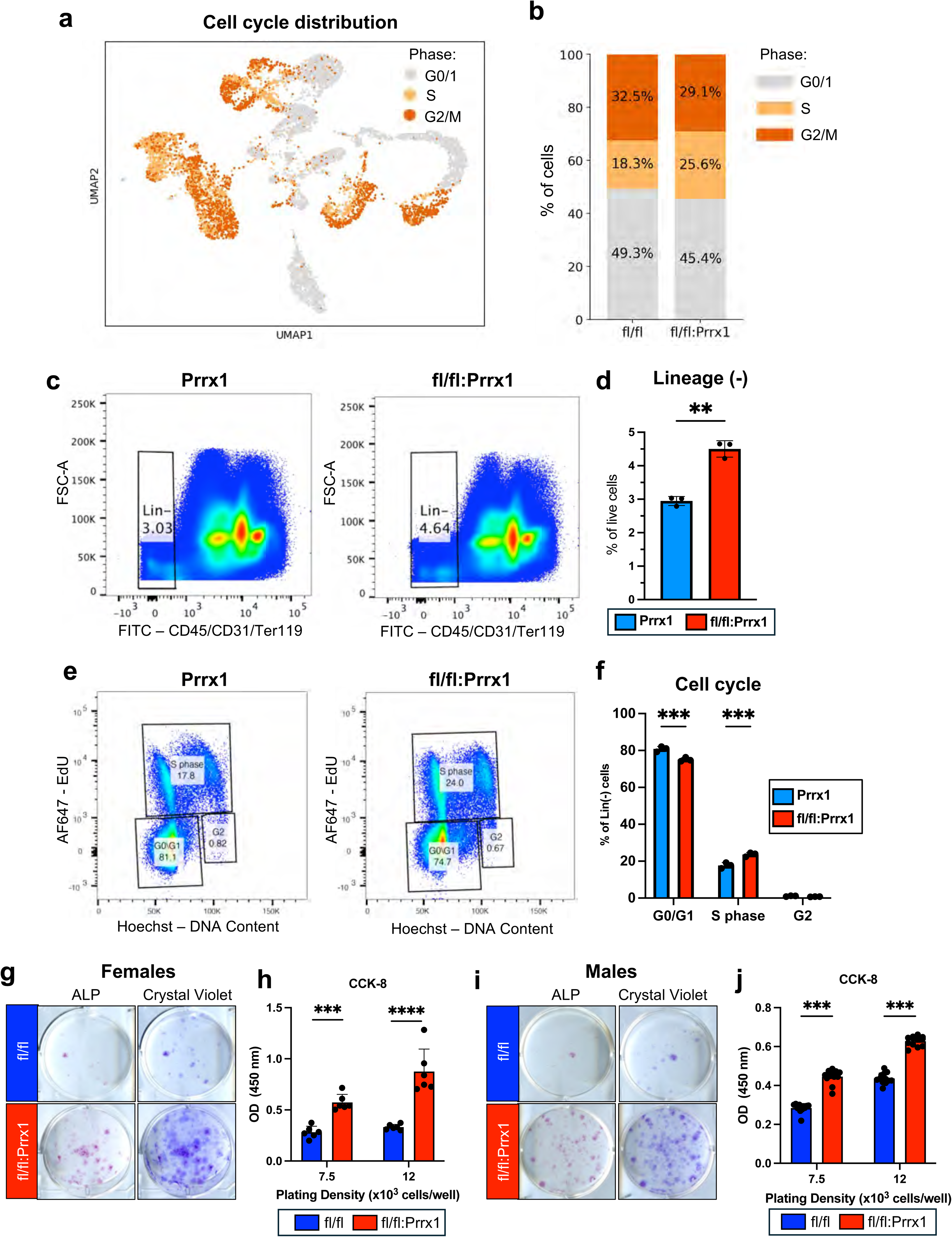
Conditional loss of *Dot1L* in Prrx1 lineage promotes expansion of Lineage-bone marrow cells. (**a**) Distribution of cycling cells across all Lineage-clusters identified by scRNAseq. (**b**) Relative proportions of G0/1, S and G2/M phase cells in Lineage-bone marrow cells from Dot1L^fl/fl^ and Dot1L^fl/fl^:Prrx1 mice. (**c**, **d)** Pseudocolor dotplots depicting the distribution of CD45^−^ CD31^−^Ter119^−^ (Lineage^−^) cells identified by flow cytometric analysis of bone marrow samples from Prrx1-Cre and Dot1L^fl/fl^:Prrx1 mice. Quantitative analysis of the relative fractions of Lineage^−^ populations in Dot1L^fl/fl^:Prrx1 mice as compared to Prrx1Cre (Dot1L sufficient) controls. n = 3 mice/genotype (**e**) *In vivo* EdU labeling confirms increased proliferative capacity of Lineage^−^ bone marrow cells in Dot1L^fl/fl^:Prrx1Cre mice. Representative plots show DNA content by Hoechst versus EdU incorporation to distinguish cell-cycle phases within the Lineage^−^ population from Prrx1-Cre controls (left) and Dot1L^fl/fl^:Prrx1 mice (right). Cells in G0/G1 were identified as Hoechst low and EdU^−^, S phase cells as EdU^+^ and G2/M cells as Hoechst high and EdU^−^. (**f**) Quantification of G0/G1, S phase and G2 cell-cycle distribution in the Lineage^−^ population for each sample. (**g, i**) *Dot1L* deletion by Prrx1Cre increased bone marrow colony forming units. Representative images of crystal violet (CV) and alkaline phosphatase activity (ALP) stained colony forming units in BMSCs from adult female and male Dot1L^fl/fl^ and Dot1L^fl/fl^:Prrx1 mice. (**h, j**) Quantitative cell proliferation in BMSCs from Dot1L^fl/fl^ and Dot1L^fl/fl^:Prrx1Cre mice from both sexes was performed using the CCK-8 kit. Cells (passage 1) were plated at two different plating densities (7.5 × 10^3^ and 12 × 10^3^ cells per well of 96 well plate). Background corrected absorbance measurements at 450nm are shown for cells from each genotype and cell seeding density. Each assay consisted of a minimum of 3 technical replicates per condition per assay, repeated 3 times. Data presented as mean ± SD; * *p* ≤ 0.05 ** *p* ≤ 0.01. *** *p* ≤ 0.001; **** *p* ≤ 0.0001.

### Dot1L deletion drives increased osteogenic and adipogenic differentiation in cultured BMSCs

Next, we interrogated whether the expansion of *Dot1L*-deficient bone marrow stromal progenitors observed *in vivo* and *in vitro* was accompanied by an altered intrinsic differentiation capacity. When stimulated with osteogenic differentiation-promoting media, cultures from Dot1L^fl/fl^:Prrx1Cre mice showed earlier mineralizing activity as compared to controls (**Figure 4a**). Quantitative analysis of Alizarin red stained cultures revealed a 3-fold increase in mineral deposition on day 10 and a 7-fold increase on day 14 of osteogenic differentiation (**Figure 4b**). Notably, even after three consecutive passages, conditions that usually diminish osteogenic potential of primary BMSCs, *Dot1L*-deficient cells continued to exhibit significantly increased Alizarin Red mineral staining (**Figure 4d**). The persistence of this phenotype across serial passaging indicates that *Dot1L* loss enforces a sustained, cell-intrinsic osteogenic differentiation capacity rather than a transient differentiation advantage. Consistent with increased mineralization, quantitative analyses of the mRNA expression of osteoblast-associated genes at day 7 of differentiation revealed a 4-fold increase in *Alp* and *Osx/Sp7, a* 5.5-fold increase *Ibsp*, and a 1.5-fold difference in *Ocn* in Dot1L^fl/fl^:Prrx1Cre versus control cells (**Figure 4c**). These observations indicate that loss of Dot1L in BMSCs leads to enhanced osteogenic commitment and mineralized matrix production in the response to osteogenic differentiation cues.

**Figure 4.**
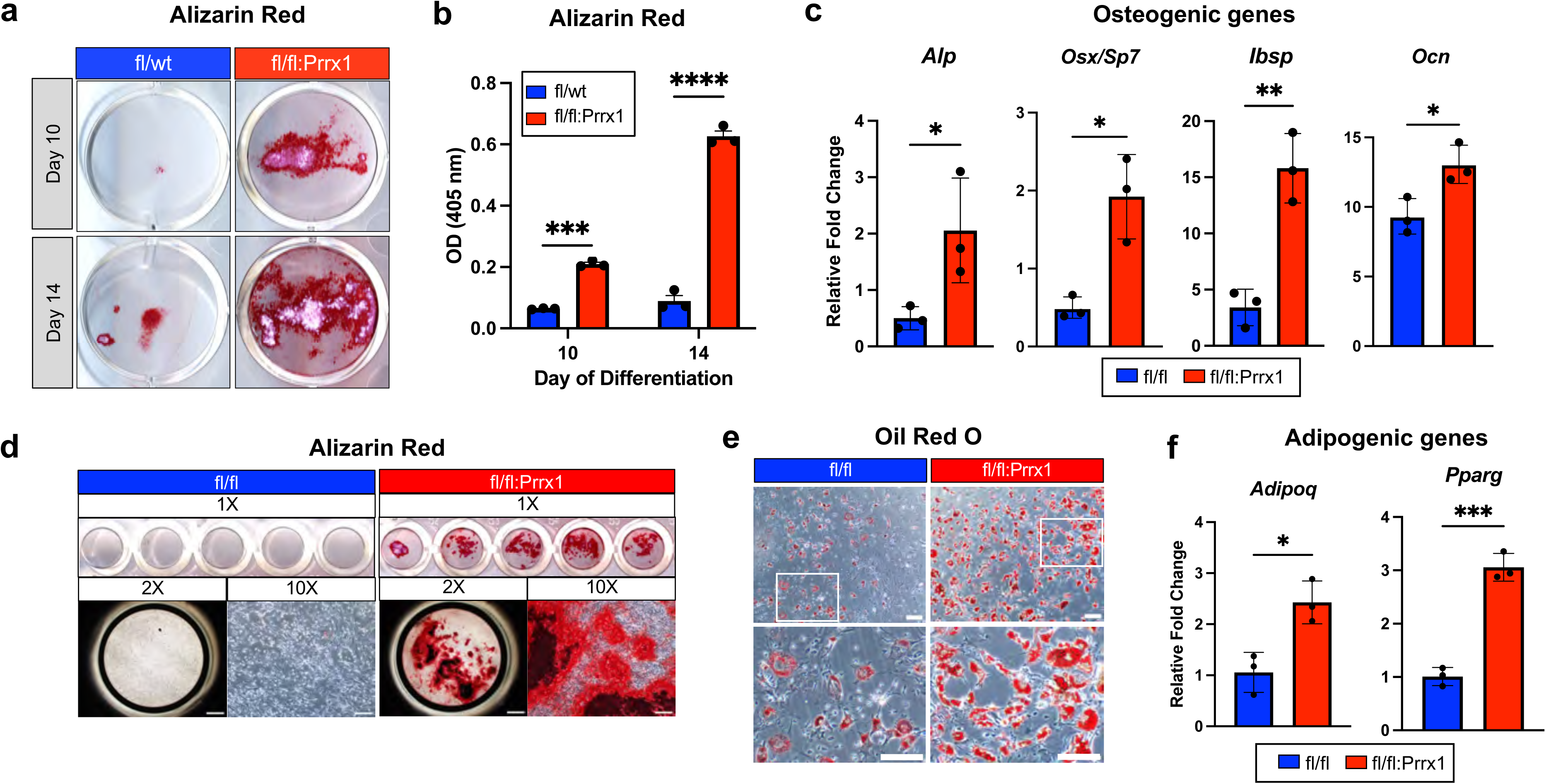
Prrx1-driven loss of *Dot1L* expression increased osteogenic and adipogenic differentiation capacity in bone marrow stromal cells. (**a**) Representative Alizarin Red staining of mineral deposits in osteogenic BMSCs from male Dot1L^fl/wt^ control and Dot1L^fl/fl^:Prrx1 mice. Cultures underwent osteogenic induction at passage 1 and were stained on days 10 and 14 of differentiation. (**b**) Quantitative analysis of Alizarin Red stain by solubilization and absorbance (405 nm). (**c**) mRNA expression of early (*Alp*), intermediate (*Osx/Sp7, Ibsp*), and late (*Ocn*) osteogenic markers on day 7 of differentiation. mRNA expression presented as relative fold change normalized to β-actin expression. (**d**) Scanned images and representative high magnification images (2x, 10x) of Alizarin Red stain depicting mineral deposition in Dot1L^fl/fl^ control cells vs Dot1L^fl/fl^:Prrx1 at day 12 of osteogenic differentiation using passage 3 cells (2x Scale bar = 1 mm; 10x Scale bar = 200 um) (**e**) Representative Oil Red O staining of lipid droplets formed within 5 days of adipogenic differentiation in BMSCs from male Dot1L^fl/wt^ control and Dot1L^fl/fl^:Prrx1 mice. Scale bar = 100 um; High magnification scale bar = 50 um. (**f**) mRNA expression of *AdiopQ* and *Pparg* at day 5 of adipogenic differentiation in Dot1L^fl/fl^:Prrx1 BMSCs relative to controls. Graphs presented as mean ± SD; * *p* ≤ 0.05 ** *p* ≤ 0.01 *** *p* ≤ 0.001; **** *p* ≤ 0.0001

In parallel studies, adipogenic differentiation assays determined that *Dot1L* loss more broadly altered stromal cell lineage commitment. Upon adipogenic induction conditions, *Dot1L*-deficient BMSCs exhibited enhanced adipocyte formation compared to controls, as evidenced by robust accumulation Oil Red O stained, lipid-filled droplets and increased adipogenic gene expression (**Figure 4e,f**). Thus, genetic removal of *Dot1L* in the Prrx1 lineage increased multilineage differentiation potential of adult BMSCs, promoting both osteogenic and adipogenic outcomes under appropriate stimulation conditions. These observations are consistent with a model in which DOT1L normally restrains stromal progenitor activation and lineage priming.

### Dot1L genetic depletion and chemical inhibition promoted the expansion of injury-responsive stromal progenitor cells

Based on robust *in vitro* and *in vivo* data, we reasoned that DOT1L normally maintains bone marrow stromal progenitors in a niche supportive state and functions as an epigenetic brake to restrain osteogenic induction. Thus, to define how DOT1L shapes SSPC responses in the early phases of the bone regenerative process we performed scRNA-sequencing of the injured marrow stromal compartment. For these studies we used an established, and highly reproducible bone marrow injury model that induces a rapid and spatially restricted intramembranous bone-forming response driven by stromal progenitor activation and proliferation, osteoblastic differentiation, and intermedullary woven bone formation.^13, 40^ Importantly, this model bypasses the formation of a cartilaginous intermediate, and therefore enables direct analysis of osteogenic lineage commitment i*n vivo*. Given the effects of Prrx1-driven *Dot1L* deletion on altered bone marrow composition and endochondral bone formation ^35^ (**Supplemental Figure 7**), we examined how haploinsufficiency influences the SSPC response to injury. Genetic haploinsufficiency in the Prrx1 lineage (Dot1L^fl/wt^:Prrx1Cre) provided a stable, lineage-restricted reduction of *Dot1L* dosage within skeletal stromal progenitors. As a key rigor element, we directly compared the cellular and transcriptional consequences of complementary genetic and pharmacologic reductions of Dot1L activity in mice (**Figure 5a**). Treatment with EPZ-5676, a highly selective, S-adenosyl-methionine (SAM) competitive inhibitor of DOT1L histone methyltransferase activity with >37,000-fold selectivity over other methyltransferases, ^41, 42^ allowed temporally controlled, acute suppression of DOT1L catalytic activity without eliciting detrimental effects on skeletal development. Effective EPZ-5676 dosing was confirmed by a significant reduction in H3K79me2 protein expression in the bone marrow of mice treated with the inhibitor for 7 days (**Supplemental Figure 6**).

**Figure 5.**
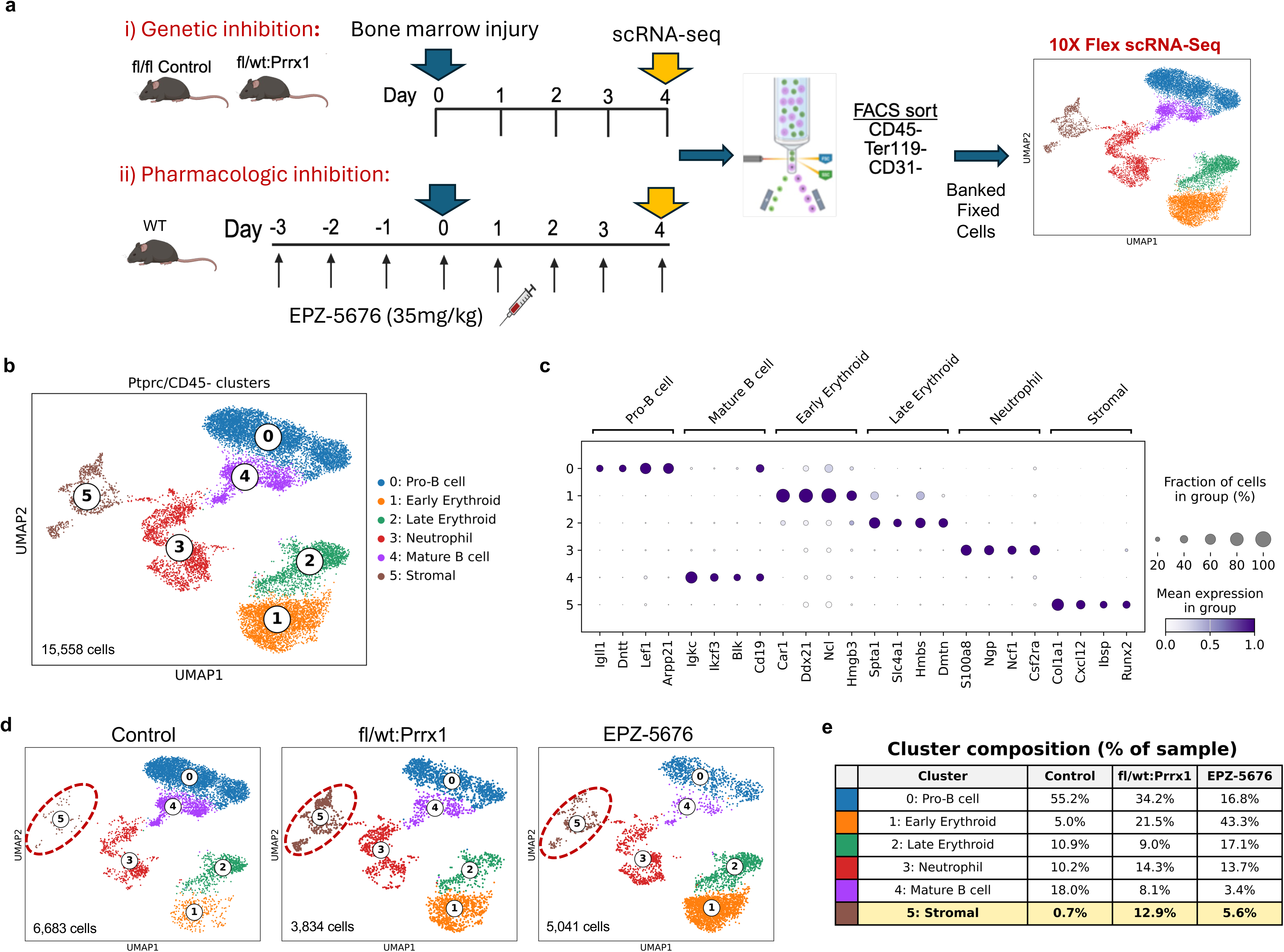
Expansion of stromal cell cluster following bone marrow injury in mice with either *Dot1L* genetic or pharmacologic perturbation. (**a**) Schematic overview of mechanical bone marrow injury model and scRNA-seq workflow. Genetic perturbation approach compared Dot1L^fl/fl^ and Dot1L^fl/wt^:Prrx1 mice. For pharmacologic inhibition, WT C57BL/6 mice were treated twice daily with 35mg/kg EPZ-5676 (i.p., 3 days prior to injury, then 4 days post-injury). Mice were sacrificed 4 days post-injury. Lineage depleted (CD45^−^Ter119^−^CD31^−^) bone marrow cells were purified and pooled (n = 3-5 mice per condition), then processed for 10x Genomics Flex scRNA-seq. (**b**) UMAP visualization of 15,558 cells following unbiased clustering and annotation based on top abundant and canonical marker genes. (**c**) Dot plot showing top discriminative marker gene sets for each cell cluster. Dot size denotes fraction of cells expressing each gene, and color scales indicate normalized mean gene expression. (**d**) UMAP projections of individual samples. The “stromal” cluster is circled. (**e**) Quantification of cell type proportions across clusters: stromal fractions are highlighted.

scRNA-seq was performed on lineage-depleted marrow cells isolated 4 days after mechanical injury, a time point chosen to capture stromal activation and lineage priming before overt mineralized bone formation.^40^ To rigorously interrogate DOT1L-dependent injury-response programs and to distinguish effects intrinsic to skeletal progenitors from those arising through acute enzymatic inhibition, we isolated Lineage^−^ cells from injured Control (Dot1L^fl/wt^ or Dot1L^fl/fl^), Dot1L^fl/wt^:Prrx1, and EPZ-5676 treated mice and processed the samples using the 10x Genomics Flex workflow (**Figure 5a**). Flow cytometric enrichment of Lineage^−^ cells was comparable across conditions (**Supplemental Figures 5a-d**), and downstream preprocessing showed similar distributions of detected genes and total UMI counts among samples after filtering for high quality cells and Harmony-based batch correction (**Supplemental Figures 5e-g**).

Integrated analysis and unsupervised clustering of 15,558 *Ptprc*^−^/CD45^−^ cells identified six transcriptionally distinct clusters representing the major stromal and hematopoietic populations present during early injury response (**Figure 5b**). Cluster identities were assigned based on canonical marker expression, and most differentially expressed and abundant genes in each cluster (**Figure 5c**). The six identified clusters included pro-B and mature B cells, early and late erythroid populations, neutrophils, and stromal cells. Comparison of UMAP plots revealed similar overall topologies among Dot1L^fl/wt^:Prrx1, EPZ-5676-treated, and Control samples. However, there were distinct differences in the relative abundance of specific cell populations among samples (**Figure 5d,e**). Quantification of cluster proportions demonstrated a reduction in lymphoid populations with concomitant expansion of erythroid cells under conditions of reduced DOT1L expression or activity (**Figure 5e**). Strikingly, the dominant DOT1L-dependent effect was observed within the stromal compartment (Cluster 5), which expanded from a rare population in Control marrow (0.7%) to a major cellular component under conditions of reduced Dot1L activity (12.9% in Dot1L^fl/wt^:Prrx1Cre; 5.6% in EPZ-5676 samples). The shared and disproportionate amplification of stromal cells across both genetic and acute pharmacologic perturbation strategies underscores DOT1L as a central epigenetic constraint on stromal progenitor activation and lineage permissiveness during early injury response.

### Genetic and pharmacologic perturbation of Dot1L elicit divergent injury-induced responses in stromal subsets

We performed high-resolution re-clustering of stromal cluster 5 to more precisely define the stromal progenitor subsets expanded by genetic dose reduction in *Dot1L* expression versus acute chemical inhibition of its catalytic activity during early injury response (**Figure 6a**). Given that the Lineage^−^ compartment comprises a broad and heterogenous spectrum of skeletal progenitor states, this approach enabled a refined resolution of SSPC subsets selectively responsive to genetic or pharmacologic perturbation of DOT1L activity. Within the stromal compartment, re-clustering resolved six transcriptionally distinct stromal sub-sets, including three populations of *Cxcl12-*abundant reticular cells (Adipo-CAR, Osteo-CAR, and Transitional CAR), Fibroblast-like mesenchymal stromal cells (Fibro-MSCs), Pericyte-like MSCs, and Chondrocytes. Consistent with the nomenclature for Cxcl12-abundant reticular subsets, the “Adipo-CAR” cluster (Cluster 3) was enriched for *Cxcl12, Lepr, Ebf3, Adipoq,* and *Foxc1*; “Osteo-CAR” cells (cluster 1) were defined by expression of *Limch1, Runx2, Alpl,* and *Ibsp*; and Transitional CAR cells (cluster 0) were identified by expression of *Basp1, Col6a1, Lox*, and *Tmsb10*. Fibro-MSCs showed selective expression of *Pdgfra*, Clec3b, *Dpt,* and *Dcn,* (**Figure 6b**), consistent with established marrow stromal identities.^1, 9, 19, 43^

**Figure 6.**
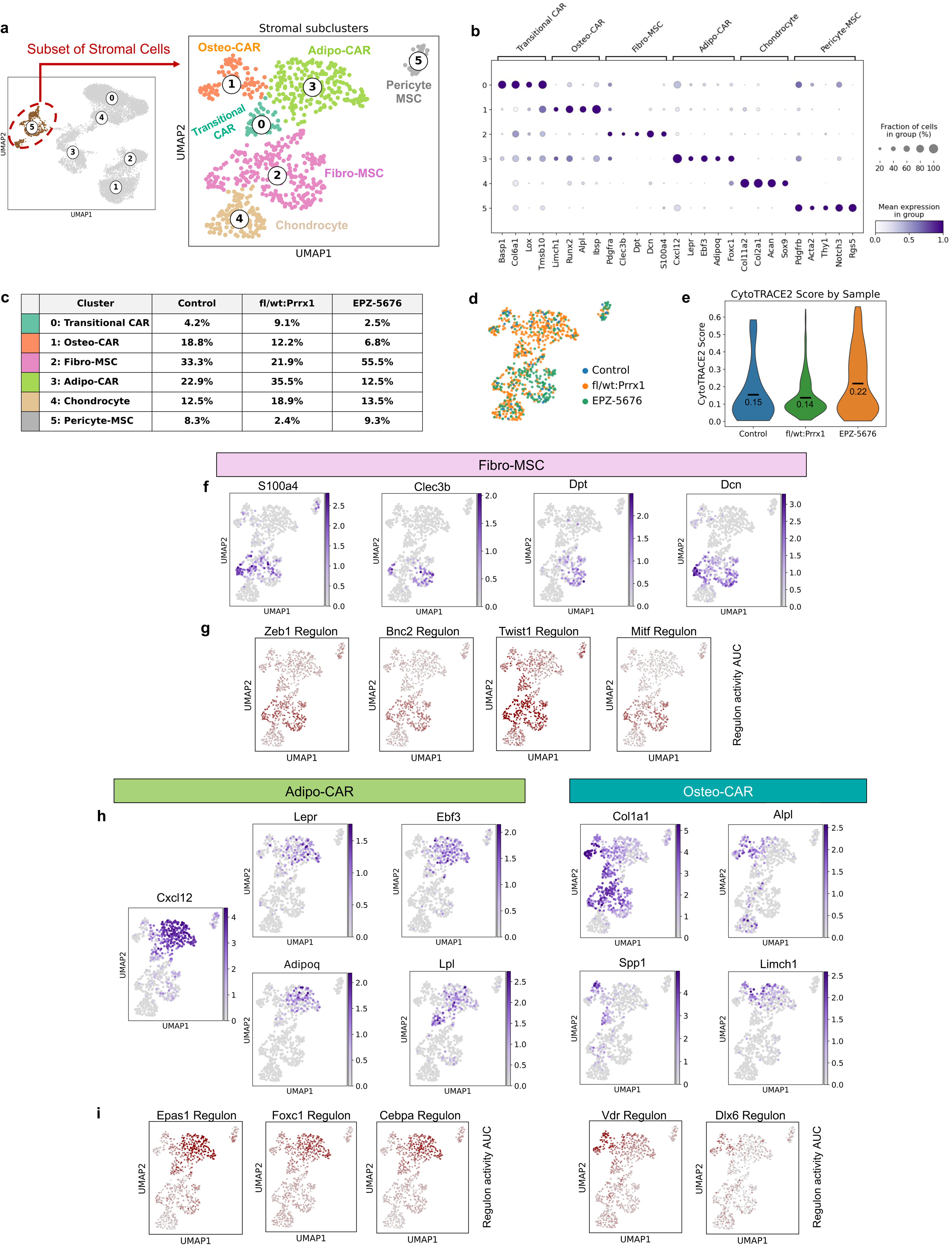
Transcriptional heterogeneity and context-dependent expansion of CAR and Fibro-MSC stromal states by Dot1L perturbation in injured bone marrow. (**a, b**) UMAP visualization of the stromal cluster that was extracted from the full dataset for secondary clustering. Dot plot compiling representative marker genes to identify cluster-specific sub-types. Dot size indicates the fraction of cells expressing each gene, and color scale represents normalized mean expression. Sub-populations included Transitional-CAR (cluster 0), Osteo-CAR (cluster 1); Fibro-MSC (cluster 2); Adipo-CAR (cluster 3), Chondrocyte (cluster 4) and Pericyte-MSC (cluster 5). **(c)** Frequency of cells within each cluster compared among Control, Dot1L^fl/wt^:Prrx1 and EPZ-5676. **(d)** UMAP projection illustrating the distribution of cells from each sample across the stromal clusters. **(e)** Violin plot depicting CytoTRACE2 stemness scores: mean score is displayed for each sample. **(f)** UMAP feature plots show expression of Fibro-MSC-associated genes (*S100a4, Clec3b, Dpt*, and *Dcn*) enriched in cluster 2 Fibro-MSC cells. **(g)** Feature plots show regulon activity of select transcription factors most active within the Fibro-MSC cluster (Zeb1, Bnc2, Twist1, and Mitf). **(h)** UMAP feature plots show expression of representative marker genes across transcriptionally distinct CAR cell sub-sets, demonstrating enrichment of Adipo-CAR genes (*Lepr, Ebf3*, *Adipoq, Lpl*), the core CAR marker *Cxcl12*, and osteogenic/ECM associated genes (*Col1a1, Alpl, Spp1, Limch1*) in Osteo-CAR cells. **(i)** Feature plots highlight the enriched regulon activity of select transcription factors within the Adipo-CAR population (Epas1, Foxc1, Cebpa) versus Osteo-CAR cells (Vdr, Dlx6).

Genetic and pharmacologic DOT1L perturbation produced distinct patterns of stromal cell expansion following injury. EPZ-5676 treatment led to preferential enrichment of the Fibro-MSC population, while genetic *Dot1L* haploinsufficiency in the Prrx1 lineage (Dot1^fl/wt^:Prrx1Cre) resulted in pronounced expansion of the three CAR subsets (**Figure 6c**). UMAP projection of sample contributions across the stromal manifold further corroborated these findings, revealing dominant enrichment of the Dot1^fl/wt^:Prrx1 cells within CAR-associated clusters and preferential localization of EPZ-5676 treated cells within the Fibro-MSC compartment (**Figure 6d**). Further, Cellular Trajectory Reconstitution Analysis using Counts expression (CytoTRACE2) revealed a notable difference in progenitor stemness state and developmental potential, with the EPZ-5676 treated sample representing an earlier injury-response program as compared to *Dot1L* genetic dose reduction (**Figure 6e**). These findings highlight a potential temporal or dose-dependent effect of genetic depletion versus pharmacologic inhibition in early marrow repair programs. To further validate the identity of the stromal populations expanded following *Dot1L* perturbation, we examined the expression patterns of canonical marker genes across stromal subclusters. UMAP feature plots showed preferential expression of ECM remodeling and fibroblast-associated genes (*S100a4, Clec3b, Dpt,* and *Dcn)* within the Fibro-MSC compartment (**Figure 6f**), consistent with previous descriptions of fibroblast-like stromal states in the bone marrow.^10^ Whereas Adipo-CAR subpopulations retained selective expression of key CAR-associated genes (*Cxcl12, Lepr, Ebf3*, *Adipoq*, and *Lpl*) and Osteo-CAR cells showed enrichment of osteogenic genes (*Col1a1, Alpl, Spp1,* and *Limch1)* (**Figures 6h**). SCENIC analysis identified distinct transcription factor regulon activity across the three CAR subsets, revealing unique regulatory programs in each stromal cell state. Fibro-MSCs exhibited higher activity of *Zeb1, Bnc2, Twist1*, and *Mitf* regulons (early mesenchymal and fibroblast transcriptional program regulators), whereas Adipo-CAR cells demonstrated increased activity of *Epas1, Foxc1*, and *Cebpa*, key regulators of marrow niche identity, and Osteo-CAR cells displayed increased *Vdr* and *Dlx5* regulon activity, consistent with osteogenic lineage commitment (**Figures 6g,i**). Together, these findings demonstrate that injury-induced stromal populations are defined by distinct transcriptional regulatory programs, and that DOT1L perturbation alters the balance of these progenitor cell states during bone reparative processes.

### Reduced Dot1L dosage reshapes CAR cell state transitions and gene regulatory programs during the early response to bone marrow injury

Given the remodeling of stromal CAR sub-populations following genetic *Dot1L* dose reduction, we next focused on the three principal CAR cell states (Transitional CAR, Osteo-CAR, and Adipo-CAR) to investigate how reduced DOT1L activity influences cell state progression during early cellular response to injury (**Figure 7a**). Diffusion pseudotime (DPT) analysis reconstructed a continuous trajectory spanning the three CAR populations, indicating progressive transitions and plasticity between stromal cell states (**Figure 7b**). Selected genes associated with DPT progression exhibited coordinated temporal changes across pseudotime (**Figure 7d**). Early pseudotime was enriched for canonical Adipo-CAR genes (ie. *Wif1, Gas6, Dpep1, Angptl1, Kitl, Pparg, Lepr, Foxc1, Esm1, Cxcl12, Ebf3, Adipoq*), intermediate peaking genes corresponded with the Transitional-CAR cell associated genes (ie. *Basp1, Angptl4, Col6a1, Igfbp7, Notch3*), whereas later pseudotime demonstrated increased expression of extracellular matrix and osteogenic genes (ie. *Dmp1, Spp1, Col1a1, Col1a2, Sparc, Spp1, Ibsp, Alpl, Runx2*), consistent with progressive acquisition of an osteogenic transcriptional program.^6, 10, 13, 44^

**Figure 7.**
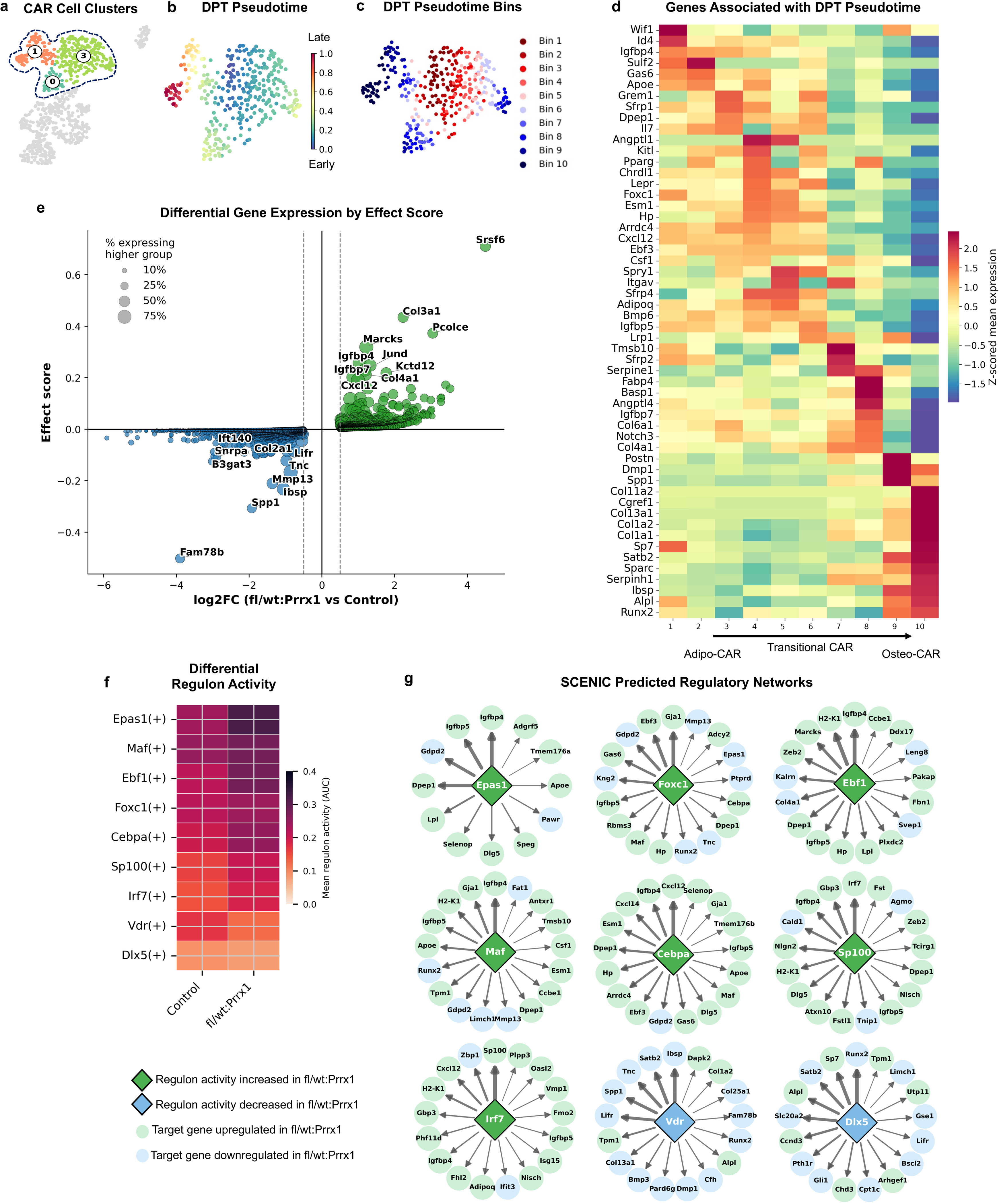
Dot1L dosage reduction alters CAR cell state transitions and regulatory networks during early marrow repair. (**a**) UMAP showing the isolation of three CAR-associated stromal populations (Adipo-CAR, Transitional CAR, and Osteo-CAR) from injured Dot1L^fl/wt^:Prrx1Cre or Control mice used for downstream trajectory and regulatory network analyses. (**b**) Diffusion pseudotime (DPT) trajectory inferred across the CAR cell continuum, with cells colored according to pseudotime progression from early to late. (**c**) Assignment of cells into ten sequential DPT pseudotime bins used for trajectory associated gene expression analyses. (**d**) Heatmap showing z-scored mean expression of pseudotime associated genes across DPT bins, illustrating the transition from Adipo-CAR to Transitional CAR to Osteo-CAR programs. (**e**) Differential gene expression between injured Dot1L^fl/wt^:Prrx1Cre and Control CAR cells, displayed as effect score versus log2 fold change. The top 10 up-regulated (green) and down-regulated (blue) genes in Dot1L^fl/wt^:Prrx1Cre CAR cells are annotated. Dot size represents the percentage of cells expressing each gene. (**f**) Mean regulon activity (AUC) for select top differentially active regulons identified by SCENIC analysis. (**g**) Predicted SCENIC regulatory networks for top differentially active regulons. Diamond nodes represent transcription factor regulons and circular nodes represent predicted target genes. Green diamonds indicate regulons with increased activity in Dot1L^fl/wt^:Prrx1Cre, whereas blue diamonds indicate decreased regulon activity. Target genes are colored according to differential expression (green-increased; blue-decreased in Dot1L^fl/wt^:Prrx1Cre vs Control). Edge thickness is proportional to inferred regulatory strength.

To determine how reduced *Dot1L* gene dosage altered these CAR cell programs, we performed differential expression analyses of the injured Dot1L^fl/wt^:Prrx1 versus Dot1L^fl/fl^ CAR cell populations. Our analysis revealed broad transcriptional remodeling associated with diminished Dot1L function, including increased expression of ECM genes (*Col3a1, Col4a1, Pcolce)* and stromal activation markers (*Marcks, Igfbp7*, *Jund*), together with reduced expression of matrix maturation and mature osteogenic genes (*Spp1, Ibsp*, *Mmp13*, and *Tnc)* (**Figure 7e**). We identified the predicted upstream regulatory mechanisms driving these transcriptional changes using SCENIC regulon analysis. The majority of regulons exhibited increased activity in Dot1L^fl/wt^:Prrx1 CAR cells relative to injured Controls (**Figure 7f**). Reconstruction of predicted regulatory networks demonstrated that these transcription factors shared common downstream targets rather than functioning as isolated regulators (**Figure 7g**). In particular, the active regulons (*Foxc1, Ebf1, Epas1, Maf*, and *Cebpa)* in stromal cells from injured Dot1L^fl/wt^:Prrx1 mice converged on a common network of CAR-associated target genes (*Ebf3, Cxcl12, Dpep1, Igfbp4/5*, and *Esm1*), whereas regulons with reduced activity (*Vdr* and *Dlx5)* regulated a distinct osteogenic transcriptional program (*Runx2, Sp7, Spp1, Alpl, Ibsp, Satb2,* and *Dmp1*). These findings suggest that reduced DOT1L dosage shifts CAR cells toward an activated stromal transcriptional state during early injury response.

### Dot1L haploinsufficiency enhances bone formation following mechanical bone marrow injury

Finally, we evaluated the functional impact of *Dot1L* genetic haploinsufficiency (Dot1L^fl/wt^:Prrx1Cre) on injury-induced intramedullary mineralization. Notably, in non-injured femurs, microCT analysis demonstrated reduced trabecular bone content in Dot1L^fl/wt^:Prrx1Cre mice versus Dot1L^fl/wt^ controls (**Supplemental Figure 7a,b**), consistent with our published data showing developmental defects in the formation and growth of the cartilage template during endochondral bone formation.^35^ Trabecular bone parameters in injured femurs from adult female Dot1L^fl/wt^:Prrx1Cre mice and Control (Dot1L^fl/fl^ or Dot1L^fl/wt^) (**Figure 8a**) revealed a robust increase in bone volume fraction (BV/TV) and trabecular number (Tb.N.), accompanied by a concomitant reduction in trabecular thickness (Tb.Th.) in injured femurs from Dot1L^fl/wt^:Prrx1 Cre mice (**Figure 8b,c**). The combined increased trabecular bone volume and trabecular number with reduced trabecular thickness denotes dense, newly formed woven bone and is consistent with accelerated regenerative bone formation following *Dot1L* dosage reduction. In parallel, calcein labeling and von Kossa staining revealed markedly increased local mineral deposition at the injury site in Dot1L haploinsufficient mice, indicative of heightened osteogenic activity and accelerated bone formation (**Figure 8d**). Injured femurs exhibited increased CXCL12 signal within the regenerated marrow space, consistent with enhanced presence or persistence of *Cxcl12* expressing stromal populations in *Dot1L* haploinsufficient mice (**Figure 8e**). In parallel, OSTERIX (Sp7) staining also revealed an upward trend in osteoprogenitor cells localized to the injury site, directly linking reduced DOT1L dosage to enhanced osteolineage engagement *in vivo*. Together, our findings indicate that *Dot1L* haploinsufficiency amplifies the early regenerative response to marrow injury, coupling expansion of injury responsive stromal and osteogenic progenitor populations with accelerated osteogenic differentiation and mineralization.

**Figure 8.**
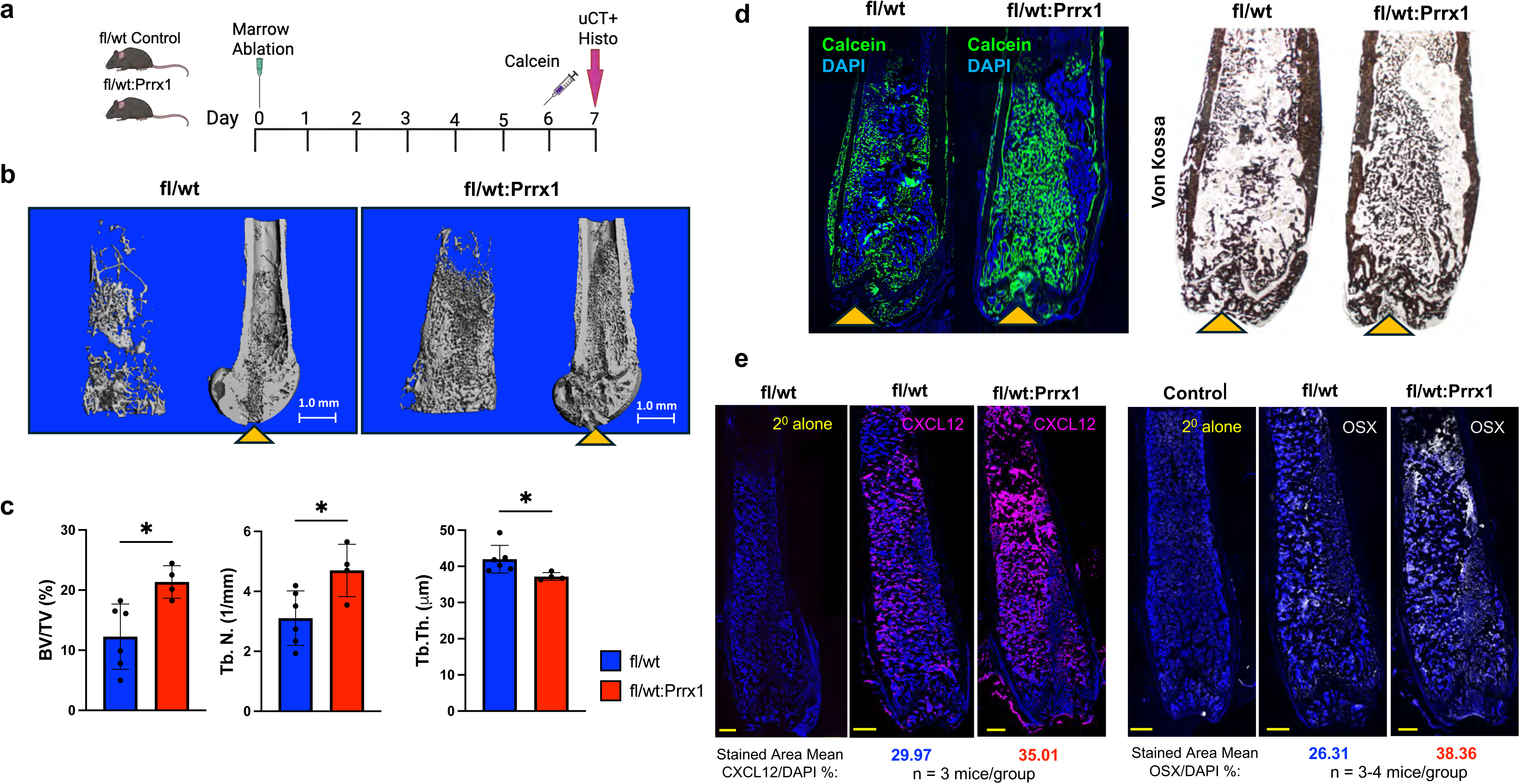
Dot1L haploinsufficiency within the Prrx1 lineage accelerates injury-induced bone regeneration. (**a**) Schematic for bone marrow injury models and phenotypic analysis of new bone formation. Mechanical bone marrow injury was induced in the left femur of skeletally mature Dot1L^fl/fl^ or Dot1L^fl/wt^:Prrx1Cre mice (day 0). Calcein mineral label was delivered by *i.p.* injection at 20 mg/kg on day 6 post-surgery, and tissue harvested on day 7. **(b)** Representative micro-CT images of trabecular cores and femur cross sections showing the path of the needle injury (yellow arrowhead) in Dot1L^fl/wt^ controls and Dot1L^fl/wt^:Prrx1 female mice on day 7 post-injury. (**c**) Quantitative trabecular bone analysis showing a significant increase in bone volume fraction (BV/TV) and trabecular number (Tb. N.), as well as a significant decrease in trabecular thickness (Tb. Th) in the regenerated bone of the Dot1L^fl/wt^:Prrx1 mice compared to controls. Graphs presented as mean ± SD; * *p* ≤ 0.05. (**d**) Representative images of Calcein mineral label (green) showing an increase in fluorescently labeled tissue in the Dot1L^fl/wt^:Prrx1 mice. Subsequent sections showing an increase in mineral deposition by Von Kossa stain with Dot1L^fl/wt^:Prrx1 compared to Dot1L^fl/wt^ control mice. Yellow arrowheads indicate path of injury**. (e)** Representative immunofluorescence images of injured femoral marrow sections from Dot1L^fl/wt^ control and Dot1L^fl/wt^:Prrx1 mice at day 7 post-injury, showing CXCL12 staining and OSTERIX (OSX) positive osteogenic progenitors within the regenerated marrow in Dot1L^fl/wt^:Prrx1 mice compared with controls. Images shown are representative sections from 3–4 independent mice per genotype. Stained area mean relative to DAPI is shown. Secondary-antibody–only controls were included for both CXCL12 and OSX immunostaining to confirm signal specificity. Scale bar = 500 um

## DISCUSSION

In this study, we identify DOT1L H3K79 methyltransferase as a key regulator of adult bone marrow stromal cell identity and lineage competence. Our findings support a model in which DOT1L maintains CAR cell identity and restrains stromal lineage induction, thereby establishing an epigenetic threshold that regulates osteogenic differentiation during regenerative processes (**Figure 9**). By integrating single cell transcriptomics, functional assays, trajectory inference, gene regulatory network analysis, and *in vivo* marrow injury models, we provide a mechanistic framework linking epigenetic regulation to CAR cell state transitions during adult bone repair. A key finding of our study indicates that DOT1L regulates the progenitor state of adult marrow stromal cells under homeostatic conditions and in response to injury, highlighting a principal role in maintaining a quiescent, niche-supportive state. Deletion of *Dot1L* augmented the intrinsic capacity of bone marrow stromal cells to undergo lineage specific differentiation, with the resulting osteogenic fate determined in a context-dependent manner.

**Figure 9.**
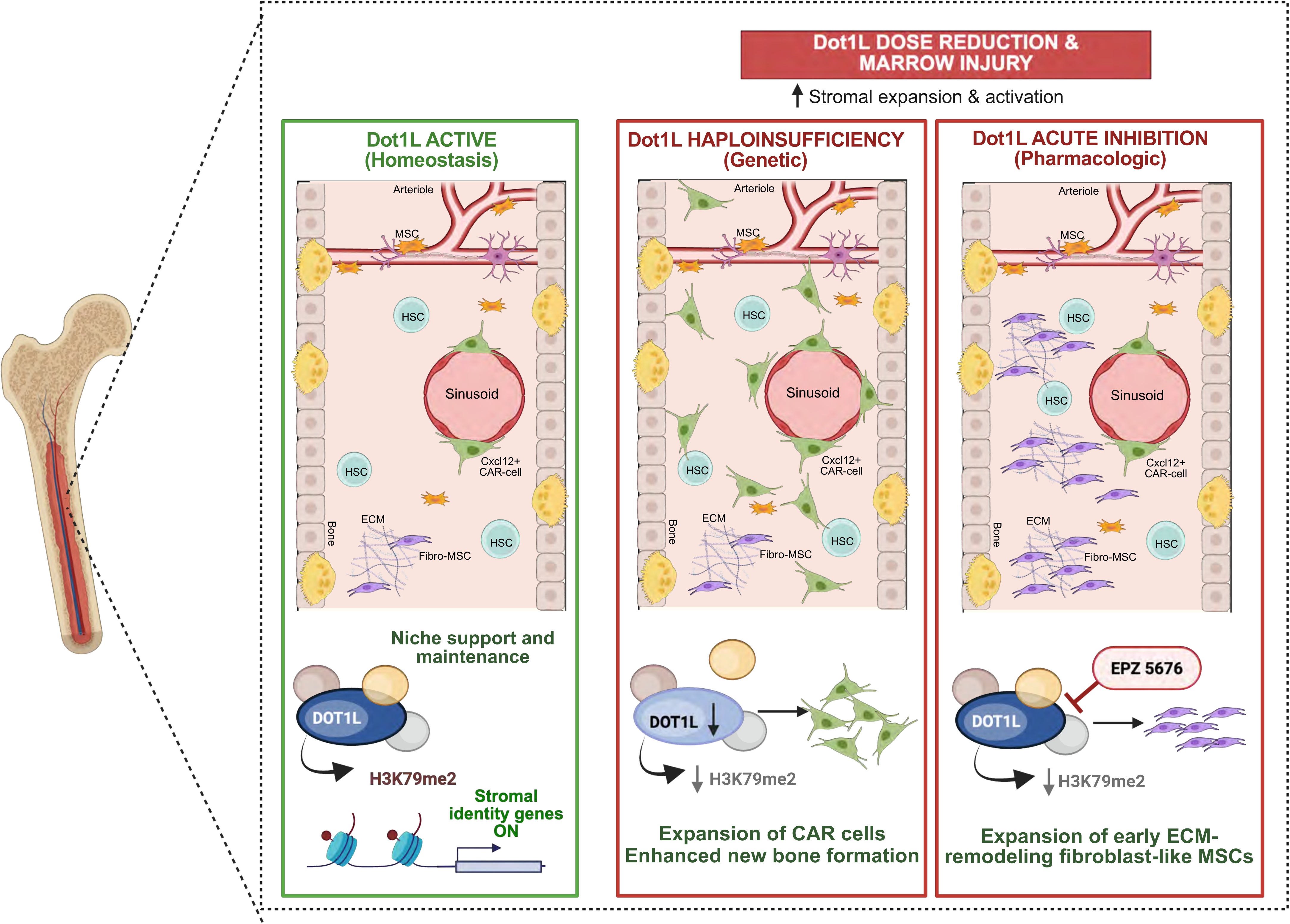
Proposed model in which DOT1L maintains niche-supportive stromal cell identity and restrains lineage induction. Under homeostatic conditions, DOT1L maintains a niche-supportive CAR cell identity and restrains transitions toward activated osteogenic stromal states. Relief of DOT1L epigenetic constraint leads to expansion of injury-responsive stromal progenitors, accelerates CAR cell state progression, and enhances regenerative bone formation following marrow injury. Schematic generated using BioRender.

Over the past several years, CAR cells have emerged as central regulators of both marrow homeostasis and skeletal regeneration.^1, 12, 13^ Originally described as specialized stromal cells supporting hematopoietic stem cells through production of CXCL12 and SCF, subsequent lineage-tracing studies demonstrated that these populations also function as osteogenic progenitors during injury repair.^1, 13, 43, 45^ More recent single-cell transcriptomic analyses further resolved substantial heterogeneity within the CAR compartment, identifying adipogenic, osteogenic, and other CAR states that occupy distinct positions along differentiation continua.^10, 11, 12, 14^ Our findings extend this framework by identifying DOT1L as an epigenetic regulator that governs the transitions between these CAR cell states. Rather than simply altering CAR cell abundance, reduced DOT1L activity shifted CAR cell composition and inferred transcription factor regulatory networks, suggesting that H3K79 methylation contributes to maintenance of CAR cell identity while simultaneously restraining progression toward injury-responsive osteogenic states.

The observation that DOT1L restrains progenitor activation represents an important conceptual advance in our understanding of the mechanisms regulating lineage permissiveness in stromal cells. Thus, rather than biasing differentiation toward a single lineage, DOT1L reduction increased multilineage differentiation capacity, suggesting that DOT1L primarily regulates progenitor state rather than lineage choice itself. These findings align with Dot1L’s established role in regulating transcriptional programs that maintain cellular identity and proliferation.^33, 46, 47^ Prior studies showed that *Dot1L* loss accelerated differentiation trajectories and disrupted the balance between proliferation and maturation in committed chondrocytes, neurons, and lymphoid cells. ^28, 34, 35, 48, 49^ In this context, the enhanced ALP⁺ colony formation and increased adipogenic differentiation observed *in vitro* here support a model in which DOT1L functions as a “gatekeeper” that regulates the timing and permissiveness of stromal cell lineage progression, while allowing environmental cues to determine ultimate lineage outcome.

A major finding of this study is the functional link between the reduced *Dot1lL* expression and activation of injury-responsive marrow stromal subsets after bone injury. Recent studies have established CAR cells as highly dynamic stromal populations that maintain the hematopoietic niche under homeostatic conditions but rapidly transition toward osteogenic states following injury ^13^. Our high resolution stromal analyses resolved distinct Adipo-CAR, Transitional CAR, and Osteo-CAR populations together with fibroblast-like stromal states, allowing reconstruction of injury-associated stromal heterogeneity. One intriguing observation was the divergent injury responses elicited by genetic and pharmacologic reduction of DOT1L activity. Although both approaches expand injury-responsive stromal populations, genetic reduction of DOT1L preferentially expanded all three CAR-associated populations, whereas acute pharmacologic inhibition produced a distinct response characterized by preferential enrichment of Fibro-MSCs and higher CytoTRACE2 scores consistent with a less differentiated injury-response program. These findings indicate that DOT1L regulates not only stromal activation but also the balance between alternative injury-responsive stromal cell states. These differences likely reflect important biological distinctions between a cumulative lineage specific reduction of DOT1L dosage and acute systemic inhibition of methyltransferase activity. Whereas genetic reduction allows stromal populations to adapt throughout adult life before injury, pharmacologic inhibition captures an immediate response to loss of catalytic activity during active repair. These findings highlight the context-dependent functions of DOT1L and suggest that both the timing and magnitude of DOT1L activity influence stromal cell state transitions during regeneration.

Our studies help reconcile apparently contrasting roles of DOT1L during skeletal development and adult tissue repair. While *Dot1L* is broadly required during embryogenesis to support progenitor proliferation and lineage progression across multiple organ systems ^29, 30, 31, 32, 33^, accumulating evidence in adult tissues ^28, 34, 50^ indicates a contrasting role in maintaining cellular identity and limiting lineage plasticity. Our previous studies demonstrated that complete deletion of *Dot1L* within embryonic limb mesenchyme disrupts skeletal development and impairs endochondral ossification, establishing an essential developmental role for DOT1L in progenitor proliferation and lineage progression.^35^ In contrast, the present study demonstrates that dose reduction of DOT1L activity in adult skeletal progenitors enhances regenerative bone formation following intramembranous marrow injury. Together, these observations support an emerging paradigm in which the function of epigenetic regulators is highly context dependent. During embryogenesis, DOT1L appears necessary to establish developmental transcriptional programs required for skeletal formation, whereas in the adult marrow it functions predominantly to maintain progenitor identity and restrain excessive activation until regenerative signals are encountered.

Several limitations in the current study should be considered. First, the Prrx1-Cre driver targets a broad limb mesenchyme-derived lineage, limiting definitive assignment of effects to specific progenitor subsets. Future studies employing inducible lineage specific Cre drivers will be important for defining the temporal and cell type specific functions of DOT1L during adult bone regeneration. Second, pooling of biological replicates for scRNA sequencing limited animal-level comparison and statistical inference, although the magnitude and consistency of effects across transcriptomic and functional analyses support biological relevance. Finally, although SCENIC identified extensive remodeling of transcriptional regulatory networks, direct chromatin profiling and assessment of H3K79 methylation occupancy will be necessary to distinguish direct DOT1L targets from secondary regulatory effects. There is growing evidence that DOT1L exerts H3K79-independent functions,^51^ thus direct catalytic inactive mutants will be required to distinguish methylation-independent mechanisms.

In summary, these studies identify DOT1L as a previously unrecognized epigenetic regulator of adult bone marrow SSPC plasticity. We propose that DOT1L maintains a niche-supportive CAR cell identity and restrains transitions toward activated osteogenic stromal states by stabilizing transcriptional regulatory networks within the adult marrow niche. By relieving a shared epigenetic constraint across multiple progenitor states, the reduction in DOT1L activity lowers the epigenetic barrier threshold, expands injury-responsive skeletal progenitors, accelerates CAR cell state progression, and enhances regenerative bone formation following marrow injury. These data establish a foundation for future studies exploring temporal modulation of DOT1L to enhance endogenous bone regeneration, while providing broader insight into how epigenetic mechanisms govern bone repair.

## MATERIALS AND METHODS

### Genetically modified mice and breeding

All breeding and genotyping strategies were followed as previously published ^35^. C57BL/6 (000664), and Prrx1Cre (005584) were obtained from The Jackson Laboratory. Female Dot1L^fl/fl^ mice originally generated and described by Bernt et al. ^52^ were crossed with Prrx1Cre male mice as we have previously described ^35^ to generate Dot1L^fl/wt^:Prrx1 mice. Dot1L^fl/wt^:Prrx1Cre male offspring were mated with Dot1L^fl/fl^ females to produce conditional knockout mice (Dot1L^fl/fl^:Prrx1Cre, designated as fl/fl:Prrx1), conditional heterozygous or haploinsufficient mice (Dot1L^fl/wt^:Prrx1Cre, designated as fl/wt:Prrx1) and wild-type littermate controls which lack expression of the Prrx1-driven Cre allele (Dot1L^fl/fl^ or Dot1L^fl/wt^). All mice were on the C57BL/6 background. Sex- and age-matched mice were used for experiments. Mice were housed on a 12-hour light-dark cycle with *ad libitum* access to food (Teklad Global 19% Protein Extruded Diet, #2918) and water under specific pathogen-free conditions at UConn Health Center for Comparative Medicine. Animal procedures were conducted in accordance with protocols approved by the Institutional Animal Care and Use Committee (IACUC) at UConn Health.

### Isolation of bone marrow stromal cell and *in vitro* proliferation assays

Total bone marrow was harvested from hindlimbs of adult (16–24-week-old) Dot1L^fl/fl^ and Dot1L^fl/fl^:Prrx1Cre mice. Marrow contents were flushed with sterile phosphate-buffered saline (PBS) using a 25-gauge needle and then filtered through a 70 µm nylon strainer. Cells were counted using an automated cell counter (Bio-Rad TC20), plated on treated tissue culture plates and maintained in growth medium consisting of DMEM supplemented with 10% fetal bovine serum and 1% penicillin– streptomycin. Cultures were maintained at 37 °C in a humidified incubator with 5% CO₂.

For colony-forming unit (CFU) assays, freshly isolated bone marrow cells from each genotype were plated at 2.12 × 10⁶ cells per well in 6-well plates and cultured for 10-11 days. Cells were fixed in 10% formalin and stained with alkaline phosphatase (ALP) solution to assess osteoblastic colony formation (CFU-Ob). Counter-staining with hematoxylin was used to visualize total fibroblastic colony formation (CFU-F). Plates were imaged using a flatbed scanner.

Cell proliferation assays were performed using the Cell Counting Kit-8 (CCK-8; Sigma-Aldrich). To enrich for adherent stromal cells and allow recovery from isolation-related stress, bone marrow stromal cells from each genotype were passaged once prior to the assay. Cells were then seeded at densities of 7.5 × 10³ or 12 × 10³ cells per well in 96-well plates and cultured for 5 days. CCK-8 reagent was added directly to the culture medium and incubated for 4.5 hours, after which absorbance was measured at 450 nm using a microplate reader (PerkinElmer EnSpire 2300). All values were background-corrected and normalized to the mean of the control group.

### Bone marrow stromal cell differentiation

Osteogenic differentiation was induced using passage 1 bone marrow stromal cells (BMSCs) seeded at 2.2 × 10⁵ cells per well in gelatin-coated 12-well plates. Upon reaching confluence, cultures were exposed to osteogenic differentiation medium consisting of growth medium supplemented with 50 µg/mL ascorbic acid and 8 µM β-glycerophosphate. Mineralized matrix deposition was assessed on days 10 and 14 of differentiation by 2% Alizarin Red staining of formalin-fixed cultures. Plates were air-dried, imaged using a flatbed scanner and Alizarin Red staining was quantified by solubilization in 10% acetic acid followed by heating at 85 °C for 10 minutes. Neutralized supernatants (10% ammonium hydroxide, pH 4.1–4.5) were analyzed by absorbance at 405 nm using a microplate reader (PerkinElmer EnSpire 2300).

In parallel, we evaluated the impact of *Dot1L* loss on adipogenic differentiation in bone marrow stromal cells. BMSCs (passage 1) were seeded at 2.5 × 10^5^ cells per well in 12-well plates. Adipogenic induction was induced once cultures reached approximately 80% confluency. Adipogenesis was induced as previously described in growth media supplemented with 3 μM rosiglitazone, 1X insulin–transferrin–selenium (ITS), and 1 μM dexamethasone ^53^. Media was replaced every 2 days. Accumulation of intracellular lipid droplets was assessed on day 5 of differentiation by Oil Red O staining of formalin-fixed cells. Oil Red O staining was visualized by brightfield microscopy.

### Quantitative PCR analysis

BMSCs undergoing osteogenic or adipogenic differentiation were washed with phosphate-buffered saline (PBS) and lysed directly in TRIzol reagent (Thermo Fisher Scientific) by scraping. Total RNA was isolated according to the manufacturer’s standard protocol, including phase separation with chloroform, RNA precipitation with isopropanol, and ethanol washes. RNA pellets were resuspended in nuclease-free water and treated with DNase I to remove genomic DNA contamination. cDNA was synthesized from purified RNA using iScript Reverse Transcription Supermix (Bio-Rad) according to the manufacturer’s instructions. The resulting cDNA was diluted to a final concentration of 5 ng/μL and used as input for quantitative PCR. qPCR was performed using Advanced Universal SYBR Green Supermix (Bio-Rad) on a CFX96 Touch Real-Time PCR Detection System (Bio-Rad). Reactions were run under standard cycling conditions with melt curve analysis to confirm product specificity. Relative mRNA expression levels were calculated using the comparative cycle threshold (Ct) method (ΔΔCt) and normalized to β-actin (*Actb*) as the internal control. Gene expression values are presented as 2^−ΔΔCt^. Murine oligonucleotide primer sequences are provided in **Table 1**.

### Mechanical bone marrow injury in mice

Bone marrow injury was performed as we have previously described ^40^. Surgeries were performed on the left femur of anesthetized female Dot1L^fl/wt^:Prrx1Cre and Cre negative (Dot1L^fl/wt^ or Dot1L^fl/fl^) controls aged 10-12 weeks. Surgeries were performed under aseptic conditions. The distal femur was accessed through the intercondylar notch, the medullary canal reamed and flushed with sterile saline until clear. The contralateral femur served as uninjured controls. Mice received extended-release buprenorphine for analgesia (Ethiqa XR, 3.25 mg/kg), and were monitored post-operatively for recovery and general health. For studies of bone formation, mice were injected with calcein mineral label on days five and six post-surgery and mice were euthanized seven days post-surgery for tissue harvest.

In a separate cohort, wild-type C57BL/6 mice were treated with EPZ-5676 (Eurofins Advinus) to achieve systemic inhibition of DOT1L methyltransferase activity. EPZ-5676 was prepared in 5% DMSO diluted in corn oil and administered via intraperitoneal injection at a dose of 35 ug/g of body weight per injection ^54^. Mice received twice-daily dosing for 4 days prior to bone marrow injury, followed by once-daily dosing for 3 days post-injury until tissue harvest.

### *In Vivo* EdU labeling and flow cytometric analysis

Adult Dot1L^fl/fl^:Prrx1Cre and Prrx1Cre male mice received a single intraperitoneal injection of 5-ethyl-2’-deoxyuridine (EdU) at 50 μg/g of body weight and maintained a 4-hour EdU labeling period to ensure adequate incorporation prior to harvest. Following euthanasia, one femur and one tibia were collected from each mouse, and the epiphyses were removed. Bone marrow was flushed with sterile PBS using a 25-gauge needle, and cell suspensions from both bones were pooled per animal. The marrow was filtered through a 70 µm nylon mesh (Nytex) to obtain a single-cell suspension. Red blood cells were lysed with ACK buffer during the permeabilization step of the Click-iT reaction. Total cell numbers were obtained using a Bio-Rad automated cell counter, and cells were aliquoted to ensure consistent staining across conditions. For each experimental sample, 10 million cells were allocated for staining, and 2-4 million cells were processed for the fluorescence-minus-one (FMO) controls, single stain, and unstained controls used for gating.

Cells were incubated with an antibody cocktail (CD45-FITC, CD31-FITC, and Ter119-FITC;1:200; **Table 2**) for 30 minutes on ice in the dark followed by washes in PBS containing 1% bovine serum albumin (BSA). Surface-labeled cells were fixed and permeabilized using the Click-iT EdU Alexa Fluor 647 Flow Cytometry Kit (Thermo Fisher) according to the manufacturer’s protocol. After two washes in the Click-iT kit buffer, fixed cells were stored overnight at 4° C in the dark. The following day, EdU incorporation was detected using Alexa Fluor 647 azide and nuclear DNA was counter stained with Hoechst 33342 (1:2000 dilution) for 30 minutes at room temperature in the dark.

Flow cytometric analysis was performed on a BD FACSymphony A5 SE analyzer (BD Biosciences). Compensation was calculated using single-stained controls and gating strategies were defined using Fluorescence Minus One (FMO) and unstained controls. Samples were acquired at low speed to minimize coincident events and at least 100,000 Lineage⁻ events were collected per sample. Post-acquisition analysis was performed using FlowJo v10.10.0 (BD Biosciences). Debris and doublets were excluded using standard forward and side scatter parameters and FSC-A/FSC-H gating. Lineage^−^ cells were identified as CD45^−^CD31^−^Ter119^−^. EdU incorporation was quantified within the Lineage^−^ compartment. Gating hierarchies were established using FMO, single-stain, and unstained controls to ensure specificity of EdU-AF647 and tdTomato signals.

### Microcomputed tomography (micro-CT)

Femurs were fixed in 10% formalin, rinsed in PBS, and stored in 70% ethanol prior to scanning. Structural analysis of day 7 post-injury femurs was performed using cone-beam x-ray micro-computed tomography (μCT40, Scanco Medical AG) with an isotropic voxel size of 8 μm (55 kVp, 145 μA, 500 ms integration). Three-dimensional reconstruction and morphometric analyses were performed using Scanco Evaluation software (v7.0). The region of interest (ROI) encompassed the intramedullary cavity of the distal femur corresponding to the injury site. A 600-slice span capturing the injured area below the growth plate was contoured manually on every 10^th^ slice to ensure consistent sampling across animals. Mineralized tissue was segmented using a global threshold (260 mg HA/cm³) applied consistently across all samples. Standard morphometric parameters, including bone volume fraction (BV/TV), trabecular number (Tb.N), and trabecular thickness (Tb.Th) were calculated using 3D direct methods.

### Histology and histochemical staining

To label newly mineralizing bone, mice were injected with calcein (i.p, 20 ug/g body weight, Sigma-Aldrich) 24 hours before harvesting. Following micro-CT imaging, femurs were processed for frozen histology to preserve fluorophore integrity. Bones were briefly rinsed in PBS, cryoprotected in 30% sucrose in PBS overnight at 4 °C, embedded in optimal cutting temperature (OCT) compound (Tissue-Tek) and frozen on dry ice. Sections (8 μm) were mounted on cryo-adhesive tape (Cryofilm 3C, Kawamoto). Calcein fluorescence with DAPI counterstain, was imaged using a Nikon Eclipse 50i or ZEISS Axioscan. To assess mineralized bone matrix, adjacent sections were stained with von Kossa by immersing in 5% silver nitrate and UV crosslinking (120 mJ/cm², twice). Brightfield images were captured using an Olympus IX71 microscope.

### Immunofluorescence staining

Immunofluorescence staining was performed on frozen bone sections collected 7 days post-injury. For OSTERIX (OSX) staining, sections were permeabilized in 0.1% Tween-20 in PBS for 15 min, followed by blocking in 10% normal goat serum (NGS) and 2% BSA in PBS for 1h at room temperature. Sections were incubated overnight at 4°C with anti-OSTERIX primary antibody (1:500; Abcam ab209484) diluted in blocking solution. After washing in PBS containing 0.1% Tween-20, sections were incubated with Alexa Fluor 647-conjugated goat anti-rabbit secondary antibody (1:300; Thermo Fisher, A21244) for 1h at room temperature, followed by additional washes prior to mounting. For CXCL12 staining, sections were rehydrated and post-fixed in 4% paraformaldehyde, followed by antigen retrieval using Proteinase K (5 μg/mL in 50 mM Tris-HCl, pH 7.5–8.0) for 5 min. Sections were permeabilized in 0.05% Triton X-100 and blocked in 10% NGS, 2% BSA, and 0.05% Triton X-100. Primary antibody incubation (anti-CXCL12, 1:100; Thermo Fisher Sci, PA5-89116) was performed overnight at 4 °C in antibody dilution buffer (PBS supplemented with 5% NGS, 1% BSA, and 0.025% Triton X-100). Following washes in PBS containing 0.05% Tween-20, sections were incubated with Alexa Fluor 647-conjugated goat anti-rabbit secondary antibody (1:300) for 1 hour at room temperature. Sections were washed and mounted with DAPI-containing glycerol mounting medium. Images were obtained using Zeiss Imager.Z1 or Zeiss Axioscan. For immunofluorescence quantitation, the distal half of the intramedullary cavity space starting below the growth plate was selected for the measured ROI. ImageJ software was used to manually threshold for staining intensity above background, the masked area was measured, and OSX/DAPI or CXCL12/DAPI area fraction was calculated.

### Re-analysis of published scRNA-seq datasets

Publicly available scRNA-seq data from CXCL12^+^ FACS-enriched mouse bone marrow stromal cells (Gene Expression Omnibus accession GSE136979) were reanalyzed in RStudio (version 2025.09.2+418) using the Seurat package v5.4.0. Raw gene expression matrices were processed following the analytical workflow described by Matsushita et al. with minor modifications for visualization and stromal cell annotation^13^. Briefly, cells were imported into Seurat, normalized using the standard Seurat workflow, and subjected to identification of highly variable genes, principal component analysis (PCA), graph-based clustering, and Uniform Manifold Approximation and Projection (UMAP) for dimensionality reduction and visualization. Cell clustering and visualization were performed using functions implemented within Seurat, following the general analytical strategy of the original study. Stromal cell populations were manually annotated based on the expression of established marker genes described in the original publication and supported by canonical markers of bone marrow stromal cell subsets. Gene expression patterns were visualized using Seurat FeaturePlot and VlnPlot functions to compare expression across annotated stromal populations. No additional statistical comparisons or differential expression analyses were performed on the published dataset; the reanalysis was used to validate and visualize expression patterns relevant to the present study.

### Visualization of publicly available bone marrow scRNA-seq data

Publicly available integrated murine bone marrow niche scRNA-seq data were explored through the Broad Institute Single Cell Portal accession SCP1248, “Resolving the Bone Marrow Niche Heterogeneity.”^39^ The resource contains 32,743 cells integrated from previously published bone marrow niche datasets using the Seurat anchor-based integration workflow, with cell populations assigned harmonized annotations based on cluster marker expression. The published UMAP coordinates and cell annotations were retained without additional computational reanalysis. Gene expression was visualized across all cells using the portal gene-expression overlay and the portal dot-plot tool. Portal-generated plots were exported and assembled without modification of the underlying data.

### scRNA-sequencing and data analysis

Bone marrow cells were harvested from femurs and tibia of Dot1L^fl/fl^ and Dot1L^fl/fl^:Prrx1Cre mice (12-20 weeks old). For injury conditions, bone marrow cells were isolated from affected femurs at 4 days post-surgery. Bone marrow stromal cell isolation and digestion were performed as previously reported (Stetsiv et al. 2025b). Cells from 4-5 mice of both sexes per genotype were pooled for a single sample for each genotype. For scRNA sequencing, cells were fixed overnight following the 10X Genomics GEM-X Flex protocol (10X). Cells were stained and sorted for CD45^−^, CD31^−^, and Ter119^−^ populations using a FACS Aria II (BD) (detailed description in Supplemental Figure 4). Library preparation and sequencing were performed by the Single Cell Biology Laboratory (SCBL) at The Jackson Laboratory for Genomic Medicine using 10X Chromium platform (5’ Gene Expression) using 5,000-10,000 cells/sample with a target depth of 365K-387K reads/cell. The GEM-X Flex-seq probe-based approach utilized the Chromium Mouse Transcriptome Probe Set v1.1.1 targeting 19,070 unique genes. Illumina base call (BCL) files were converted to FASTQs using bcl2fastq v2.20.0.422 (Illumina) and trimmed to a 28-10-10-90 asymmetric read configuration. The resulting FASTQ files were processed using the Cell Ranger multi pipeline (10x Genomics, v9.0.1). Reads were aligned to the GRCm39 probe set reference, and the probe barcodes were used to demultiplex the pooled samples into individual datasets. Filtered feature barcode matrix files were used for downstream clustering and analysis using Scanpy v1.11.4. Ambient RNA was removed using SoupX (v1.6.2), and doublets were identified using scDblFinder (v1.23.4). Batch correction was performed using HarmonyPy (v0.0.10). Genes present in fewer than 10 cells were removed. Low quality cells were excluded if they expressed fewer than 200 genes or deviated by more than 5 median absolution deviations for metrics including gene counts, total counts, proportion of top 20 expressed genes, and mitochondrial, ribosomal, and hemoglobin content (Supplemental Figure 5). Data were transformed using the shifted logarithm for clustering and differential expression analysis (log2fold change). CD45-enrichment efficiency was validated by *Ptprc* expression and only cells with zero *Ptprc* expression were subset for subsequent analyses. Cells were clustered with the Leiden algorithm based on the top 4,000 variable genes (Seurat v3 method), using 20 principal components neighbor computation. Cell clusters were annotated by abundant marker and known gene expression.

To identify biologically meaningful transcriptional differences between experimental groups, differential gene expression was quantified using an effect size-based metric rather than relying solely on statistical significance. For each gene, log-transformed normalized expression, the difference in the proportion of expressing cells, and the difference in mean expression were calculated between experimental groups. An integrated effect score was computed to summarize the magnitude and consistency of transcriptional change while accounting for both expression level and cellular prevalence. Genes exhibiting large changes in both average expression and the fraction of expressing cells received higher effect scores than genes showing isolated changes in a single metric. For visualization and downstream analyses, genes were filtered to retain those expressed in at least 5% of cells in either comparison group and exhibiting an absolute log2 fold change ≥0.5. Effect scores were used to rank genes for pathway enrichment analyses and to prioritize biologically relevant transcriptional changes.

Genes were ranked according to their effect scores and analyzed by pre-ranked Gene Set Enrichment Analysis (GSEA) using GSEApy. Enrichment analyses were performed using Gene Ontology Biological Process 2026. Gene sets containing fewer than five or more than 500 genes were excluded. Enrichment significance was estimated using 1,000 phenotype permutations, and normalized enrichment scores (NES) were used to compare pathway activity across experimental groups.

Cell cycle phase scores were calculated using the Scanpy score_genes_cell_cycle function with the canonical S-phase and G2/M-phase marker genes as previously described.^55^ Cells were assigned predicted G1, S, or G2/M phases based on the relative S and G2/M module scores. Cell-cycle scores were used for visualization and comparison of proliferative states between experimental groups but were not regressed during preprocessing.

Pseudotime trajectories for the three CAR populations were inferred using Scanpy diffusion pseudotime (DPT). Neighborhood graphs were recomputed from the stromal subset using 30 principal components and 15 nearest neighbors. The root cell was selected using the highest CytoTRACE2 score within the Adipo-CAR cluster. Unbiased and known CAR-associated genes exhibiting dynamic expression across pseudotime were identified by calculating average expression within 12 equally spaced pseudotime bins and visualized as a heatmap.

Gene regulatory networks were inferred using RegDiffusion followed by pySCENIC. RegDiffusion was used to infer transcription factor-target relationships from normalized stromal cell expression matrices after removal of zero-variance genes and genes expressed in less than 5 cells. The resulting transcription factor-target interaction network was filtered to retain the highest-confidence targets for each transcription factor and used as input for pySCENIC. Motif enrichment analysis was performed using the mm10 cisTarget motif databases, and regulon activity scores were calculated using AUCell. Differential regulon activity between experimental groups was quantified as differences in mean AUCell scores. Candidate regulatory interactions were integrated with differential expression analyses to identify transcription factors associated with altered stromal cell states following genetic reduction or pharmacologic inhibition of DOT1L.

### Statistical Analyses

Statistical analyses were performed using GraphPad Prism (Version 10.5.0). All quantitative data are presented as mean ± standard deviation (SD), unless otherwise indicated. Statistical comparisons between two groups were performed using Student’s *t*-tests. Comparisons involving multiple groups were analyzed by one-way or two-way analysis of variance (ANOVA), followed by Tukey’s post hoc multiple-comparison tests, as appropriate. For scRNA-seq analyses, differential gene expression was assessed by Wilcoxon ranking, using log-transformed data and appropriate statistical thresholds as described in the Methods.

---

**Supplemental Figure 1.**
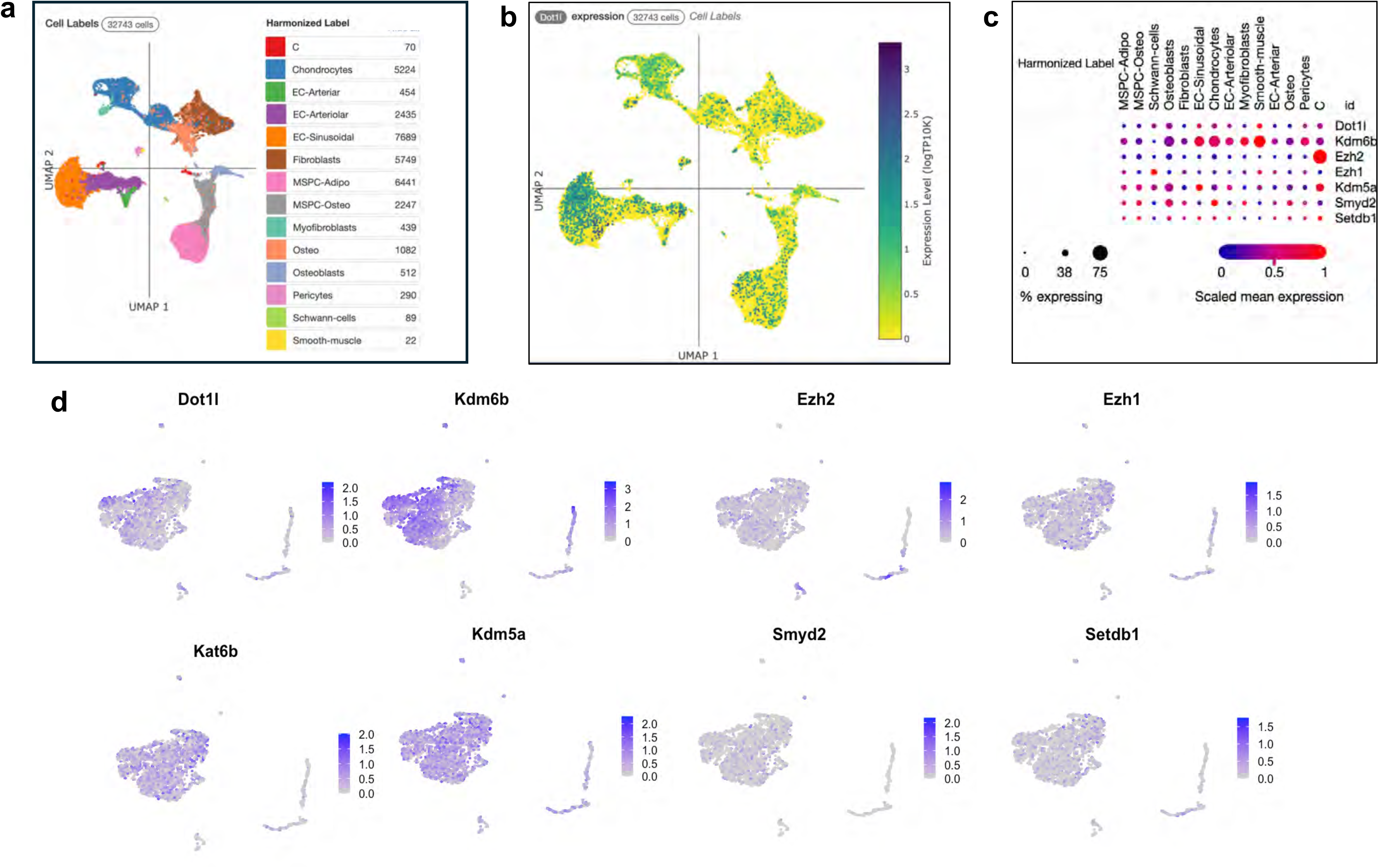
D*o*t1L is part of a broad network of chromatin modifying epigenetic regulators within marrow stromal populations. **(a)** Integrated scRNA-sequencing analysis of murine bone marrow cells compiled across multiple experimental and physiological conditions (Broad Single Cell Portal, study SCP1248)^39^. UMAP representation of transcriptionally defined clusters annotated based on established lineage and niche marker expression, including stromal, hematopoietic, endothelial, and other niche-associated populations. **(b)** Feature plot showing *Dot1L* expression projected onto the clustered dataset, demonstrating broad transcript detection across marrow populations with consistent expression within multiple stromal clusters. (**c)** Dot plot summarizing scaled expression levels (color scale) and the fraction of cells expressing *Dot1L* alongside select histone modifying enzymes and chromatin associated regulators across stromal clusters, situating DOT1L within a broader epigenetic regulatory landscape in bone marrow stromal cells. **(d)** Expression of epigenetic regulators across CXCL12^+^ bone marrow niches. UMAP feature plots showing expression patterns of selected chromatin modifying enzymes associated with regulation of osteogenic differentiation across single-cell transcriptomes from fluorescence-activated cell sorted CXCL12^+^ bone marrow stromal cells (GSE136979)^13^.

**Supplemental Figure 2.**
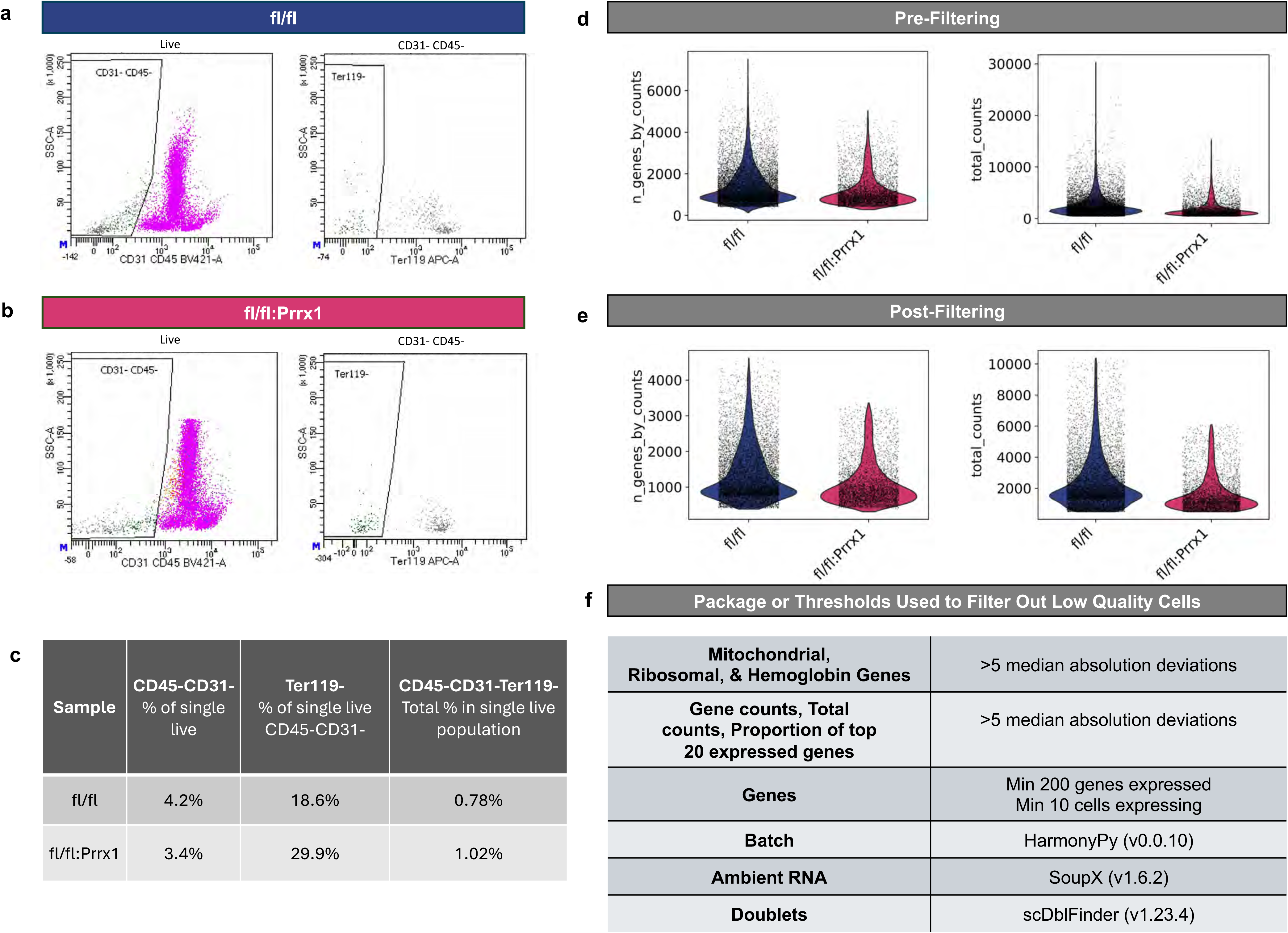
FACS lineage-cell enrichment and quality control filtering criteria for scRNA sequencing of Dot1L^fl/fl^ and Dot1L^fl/fl^:Prrx1Cre cells. **(a, b)** Representative flow cytometry plots showing sequential gating to enrich for Lineage^−^ stromal cells (CD45^−^, CD31^−^, Ter119^−^) from Dot1L^fl/fl^ control and Dot1L^fl/fl^:Prrx1 samples used for single-cell RNA sequencing. Initial exclusion of hematopoietic (CD45^+^) and endothelial (CD31^+^) populations was followed by removal of erythroid lineage cells (Ter119^+^). **(c)** Table summarizing the percentage of cells within each gated population for each sample. **(d, e)** Violin plots showing the distribution of detected genes per cell and total UMI counts per cell across Dot1L^fl/fl^ control and Dot1L^fl/wt^:Prrx1 samples prior to and following initial pre-processing. Overlaid points represent individual cells. **(f)** Table summarizing quality control filtering criteria applied prior to downstream analysis, including removal of cells with high mitochondrial (mt) or ribosomal/hemoglobin (Ribo, Hb) gene content; gene counts per cell, total UMI counts, and the proportion of top 20 expressed genes; gene inclusion thresholds (minimum 200 genes per cell and genes expressed in at least 10 cells); batch correction; and removal of ambient RNA and cell doublets.

**Supplemental Figure 3.**
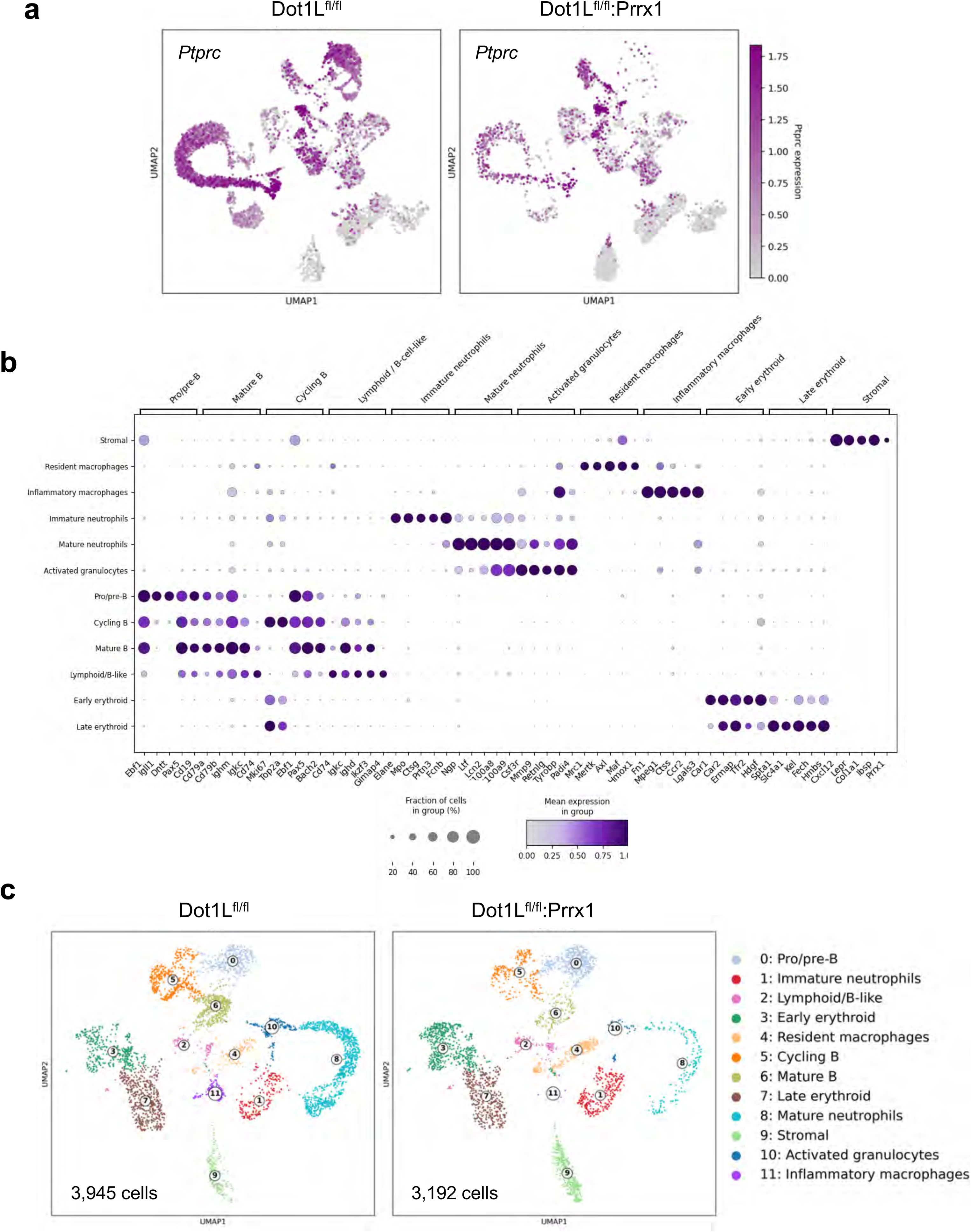
scRNA-seq cluster annotation of intact Dot1L^fl/fl^ control and Dot1L^fl/fl^:Prrx1Cre cells. **(a)** UMAP showing residual *Ptprc* expression across samples. Cells lacking *Ptprc* (CD45) expression were used for downstream clustering and analyses. **(b)** Dotplot shows mean expression and fraction of cells expressing the top differential genes and canonical markers used for cluster annotation. **(c)** UMAP showing 3,945 Dot1L^fl/fl^ and 3,192 Dot1L^fl/fl^:Prrx1Cre cells that clustered into 12 transcriptionally distinct clusters, represented by lymphoid B cells (clusters 0, 2, 5, 6), granulocytes/neutrophils (clusters 1, 8, 10), erythroid cells (clusters 3, 7), macrophages (clusters 4, 11) and stromal cells (cluster 9).

**Supplemental Figure 4.**
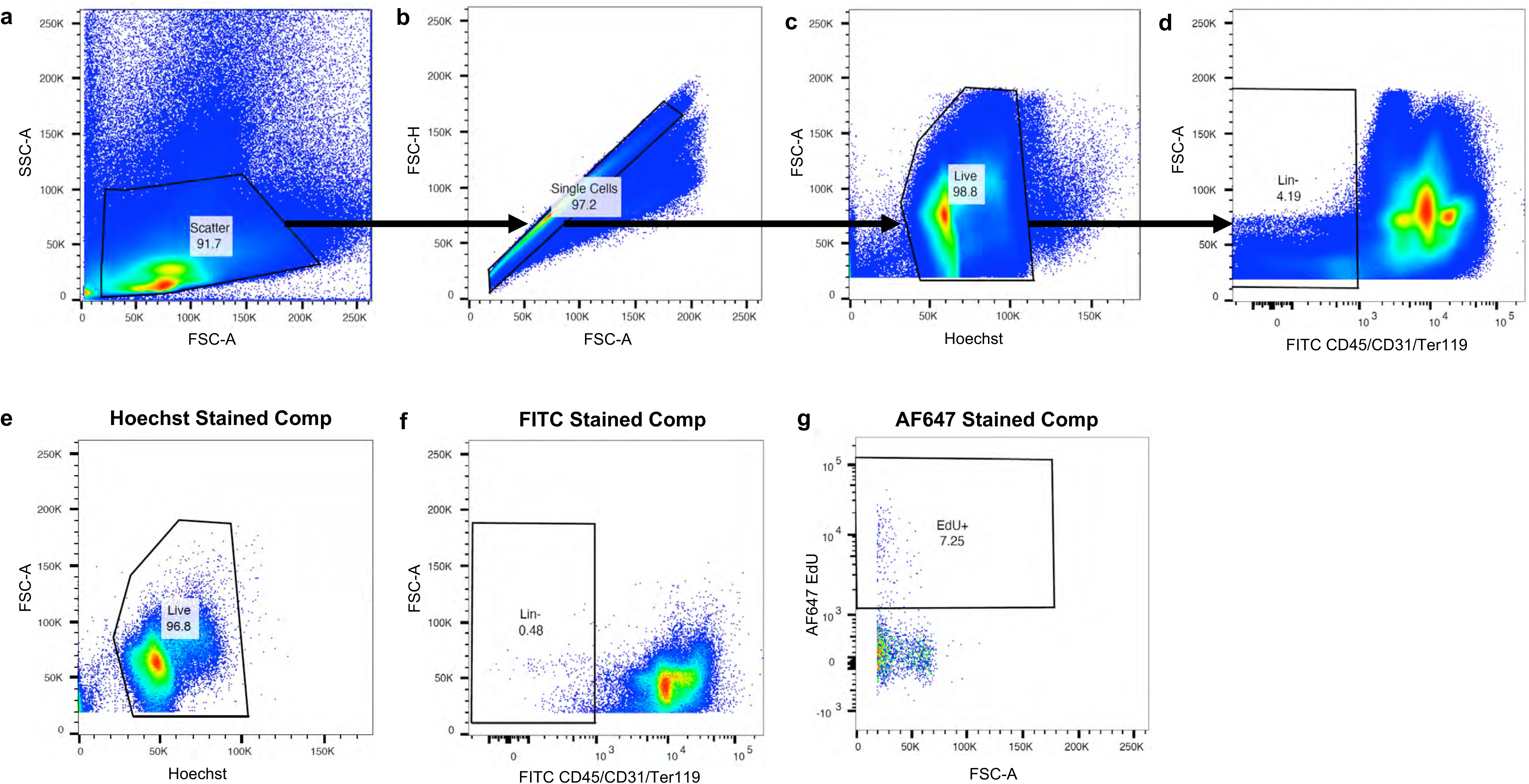
Flow cytometry gating strategy and controls for *in vivo* proliferation analysis. Sequential gating strategy used to identify Lineage^−^ (CD45^−^CD31^−^Ter119^−^) bone marrow stromal cells for EdU proliferation analysis. **(a)** Debris were excluded by first gating on forward scatter (FSC-A) versus side scatter (SSC-A). **(b)** Singlets were isolated by gating on FSC-H versus FSC-A, **(c)** followed by selection of nucleated cells based on Hoechst fluorescence. **(d)** Lineage^−^ stromal cells were identified by excluding FITC-labeled CD45^+^, CD31^+^, and Ter119^+^ hematopoietic and endothelial populations. **(e)** Hoechst only stained compensation control **(f)** FITC-only stained compensation control (CD45/CD31/Ter119 panel). **(g)** AF647-only stained compensation control (EdU^+^). All gating thresholds were determined using single color compensation, FMO, and unstained controls to ensure accurate identification of viable Lineage^−^ and EdU^+^ populations.

**Supplemental Figure 5.**
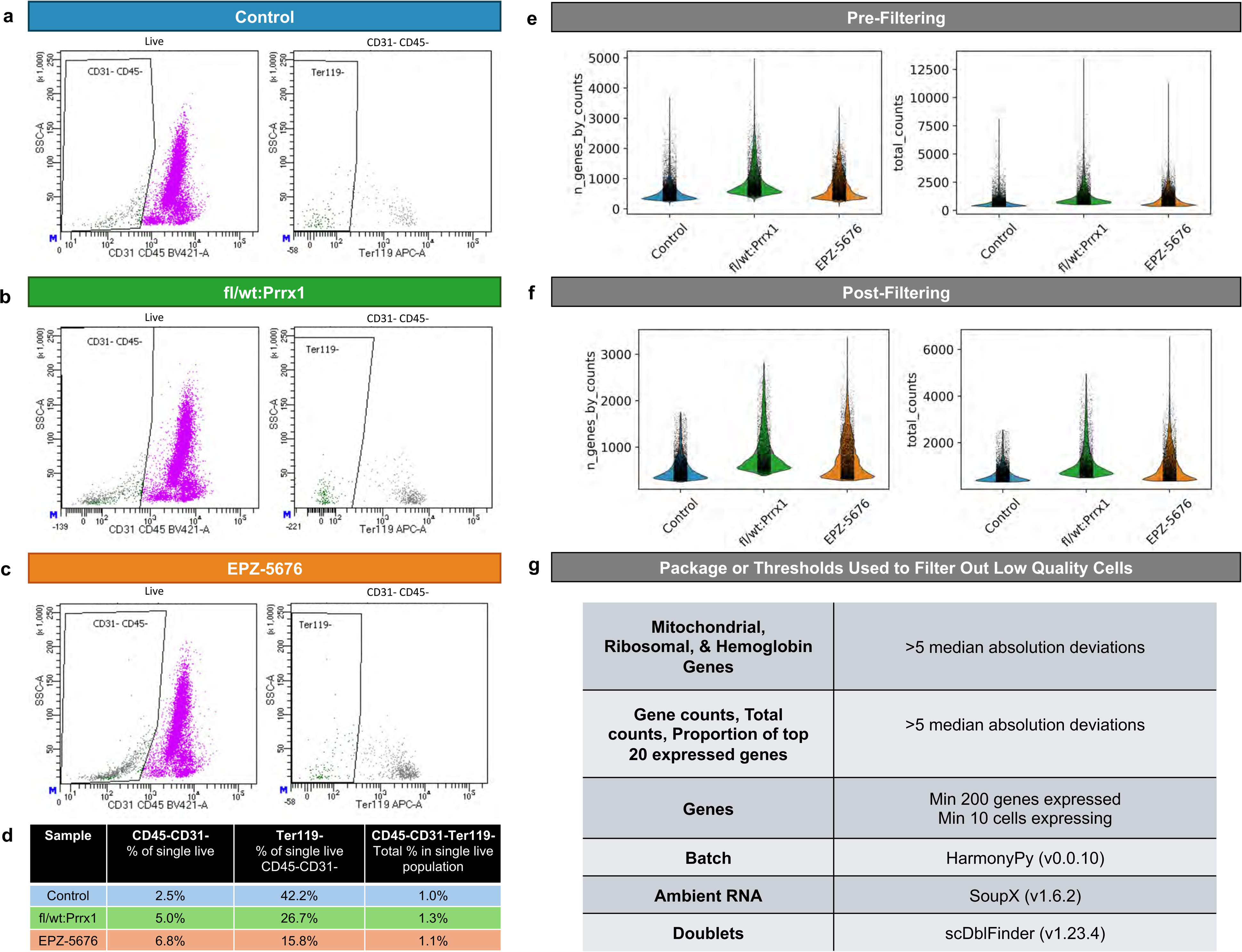
FACS Lineage^−^ cell enrichment and quality control filtering criteria for scRNA-seq of D4 post-injury Control, Dot1L^fl/wt^:Prrx1Cre, and EPZ-5676 treated cells. **(a, b, c)** Representative flow cytometry plots showing sequential gating to enrich for Lineage^−^ stromal cells (CD45^−^, CD31^−^, Ter119^−^) from 4-day post marrow injury Control (Dot1L^fl/fl^ or Dot1L^fl/wt^), Dot1L^fl/fl^:Prrx1, or EPZ-5676 treated mice used for single-cell RNA sequencing. Initial exclusion of hematopoietic (CD45^+^) and endothelial (CD31^+^) populations was followed by removal of erythroid lineage cells (Ter119^+^). **(d)** Table summarizing the percentage of cells within each gated population for each sample. **(e, f)** Violin plots showing the distribution of detected genes per cell and total UMI counts per cell across Dot1L^fl/fl^ control and Dot1L^fl/wt^:Prrx1 samples prior to and following initial pre-processing. Overlaid points represent individual cells. **(g)** Table summarizing quality control filtering criteria applied prior to downstream analysis, including removal of cells with high mitochondrial (mt) or ribosomal/hemoglobin (Ribo, Hb) gene content; gene counts per cell, total UMI counts, and the proportion of top 20 expressed genes; gene inclusion thresholds (minimum 200 genes per cell and genes expressed in at least 10 cells); batch correction; and removal of ambient RNA and cell doublets.

**Supplemental Figure 6.**
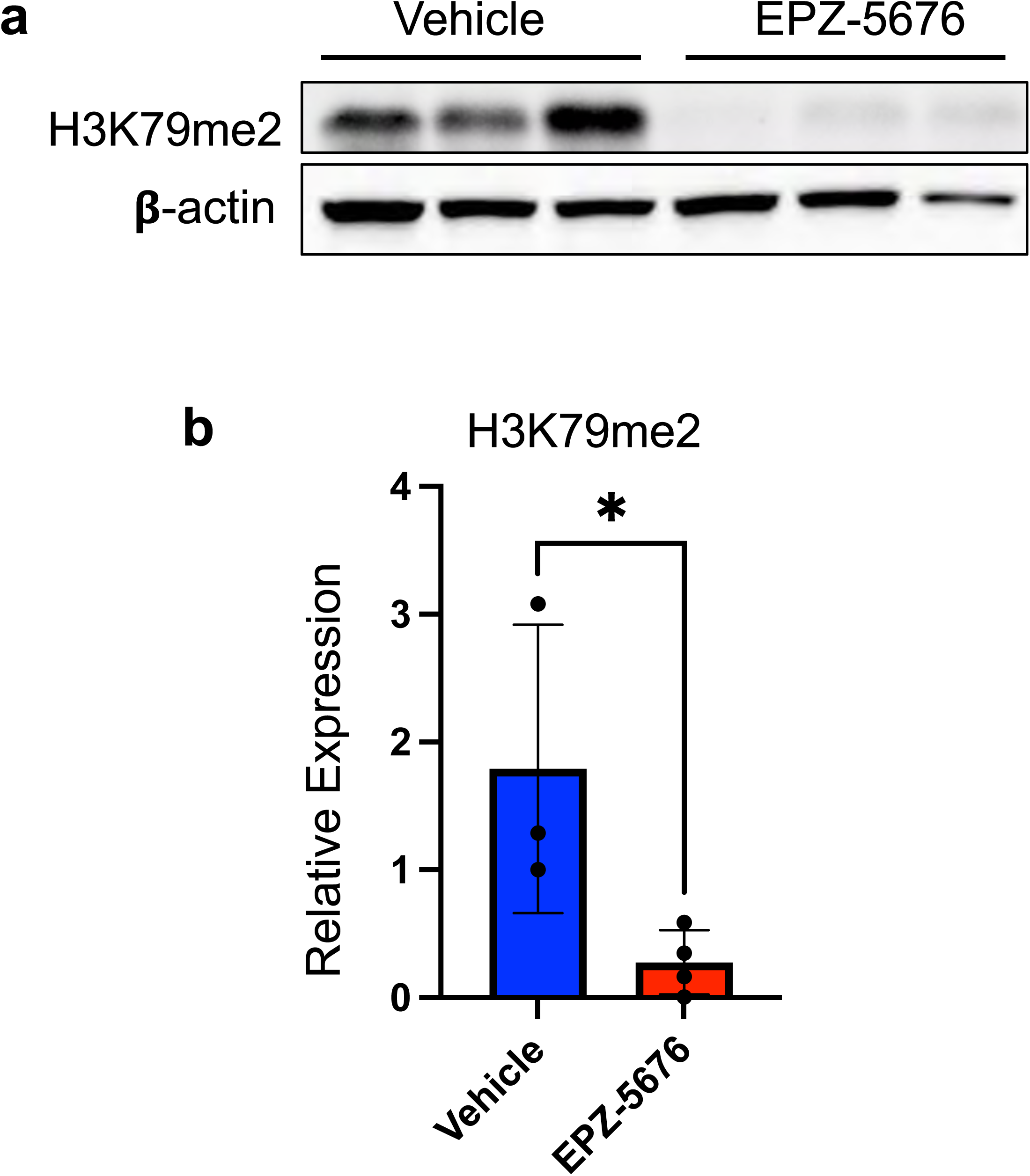
Validation of DOT1L inhibitor dosing and efficacy *in vivo*. (**a**) Representative western blot analyses of bone marrow cells from vehicle and EPZ-5676-treated mice. C57BL/6 mice were treated with the DOT1L inhibitor EPZ-5676 (35 mg/kg, twice daily) or vehicle (5% DMSO in corn oil) for 7 consecutive days. Bone marrow cells were harvested, subjected to red blood cell lysis, and analyzed by western blot for H3K79me2 protein expression. (**b**) Quantification of H3K79me2 signal normalized to *β*-actin. Data are presented as mean ± SD; * *p* < 0.05 by two-tailed unpaired t-test. n = 3 mice/group.

**Supplemental Figure 7.**
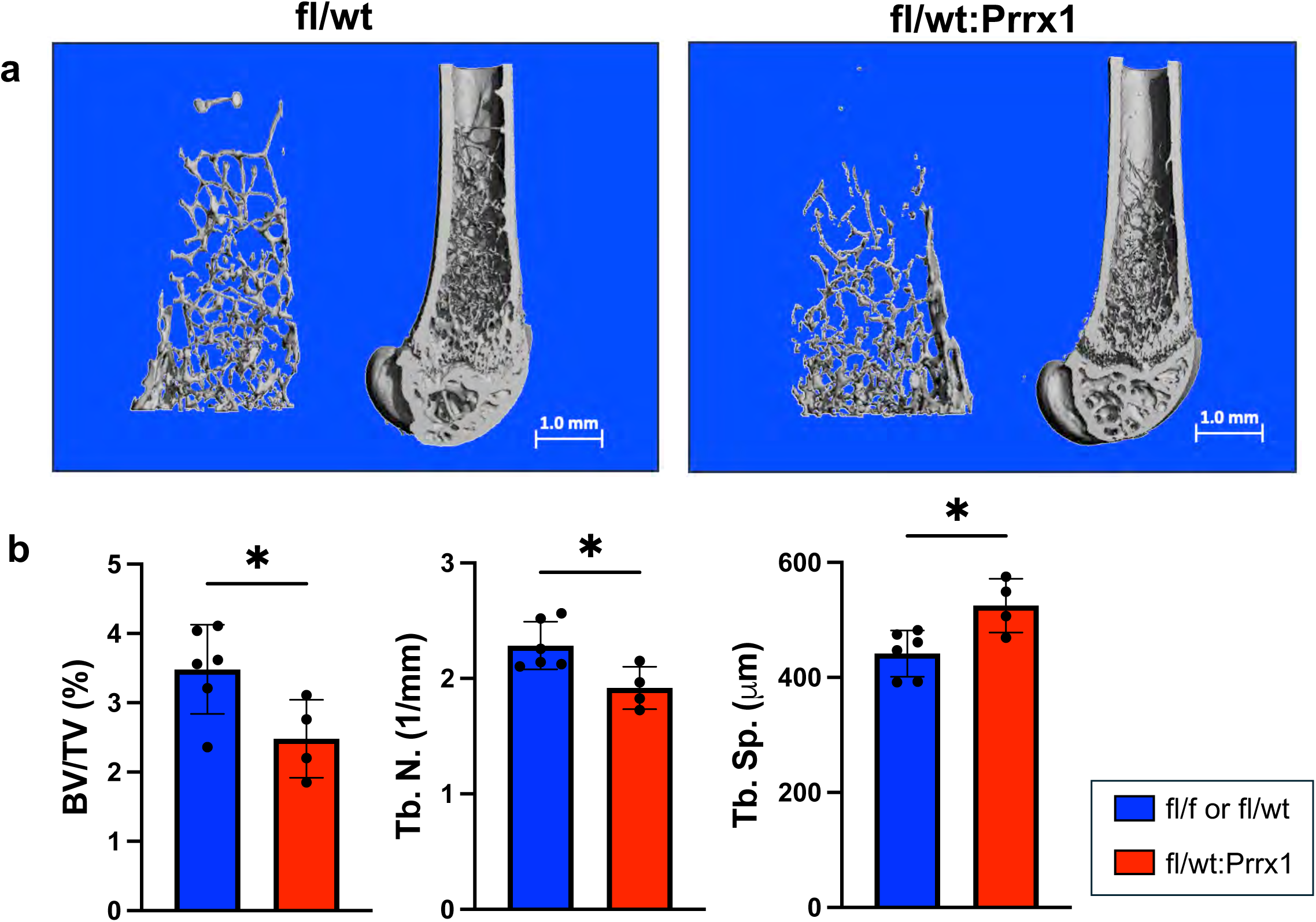
D*o*t1L haploinsufficiency leads to a reduction in bone volume at baseline in female mice. (**a**) Representative microCT images of cross sections and trabecular cores of intact femurs from Dot1L^fl/wt^ Control and Dot1L^fl/fl^:Prrx1 female mice. (**b**) Quantitative trabecular bone analysis showed a minor but significantly reduced bone volume fraction (BV/TV) and trabecular number (Tb. N.), and a concomitant increase in trabecular spacing (Tb. Sp.) in contralateral (non-injured) femurs from female Dot1L^fl/wt^:Prrx1 mice compared to controls. Graphs presented as mean ± SD; * *p* ≤ 0.05.

## ACKNOWLEDGEMENTS

We would like to thank Dr. Evan Jellison of the UConn Health Flow Cytometry Core, and Renata Rydzik and Dr. Benjamin Sinder of the UConn MicroCT Facility. We thank Ms. Marisha Pinto for technical assistance. We gratefully acknowledge the contributions of the Single Cell Biology Lab and Genome Technologies services at The Jackson Laboratory for expert assistance with the work described herein. These shared services are supported in part by the JAX Cancer Center (P30 CA034196).

## AUTHOR CONTRIBUTIONS

**Marta Stetsiv**: Writing – original draft, review and editing. Investigation, visualization, formal analysis, data curation and conceptualization. **Sakinah Abdualsalm**: Investigation, formal analysis and data curation. **Alex Tress**: Investigation. **Drew Dauphinee**: Investigation, formal analysis and data curation. **Kerry Cobb**: Formal analysis and supervision. **Archana Sanjay**: Writing – review and editing, supervision, investigation, funding acquisition and conceptualization. **Rosa Guzzo**: Writing – review and editing, supervision, project administration, investigation, funding acquisition and conceptualization.

## MATERIALS AND CORRESPONDENCE

Further information and requests for resources and reagents should be directed to and will be fulfilled by Rosa Guzzo and Archana Sanjay.

